# A CLN6-CLN8 complex recruits lysosomal enzymes at the ER for Golgi transfer

**DOI:** 10.1101/773804

**Authors:** Lakshya Bajaj, Alberto di Ronza, Pengcheng Zheng, Aiden Eblimit, Rituraj Pal, Jaiprakash Sharma, Dany Roman, John R. Collette, Richard N. Sifers, Sung Y. Jung, Rui Chen, Randy W. Schekman, Marco Sardiello

## Abstract

Lysosomal enzymes are synthesized in the endoplasmic reticulum (ER) and transferred to the Golgi complex by interaction with the Batten disease protein CLN8. Here we investigated the relationship of this pathway with CLN6, an ER-associated protein of unknown function that is defective in a different Batten disease subtype. Experiments focused on protein interaction and trafficking identified CLN6 as an obligate component of a CLN6-CLN8 complex (herein referred to as EGRESS: <u>E</u>R-to-<u>G</u>olgi relaying of <u>e</u>nzymes of the ly<u>s</u>osomal <u>s</u>ystem) which recruits lysosomal enzymes at the ER to promote their Golgi transfer. Simultaneous deficiency of CLN6 and CLN8 did not aggravate mouse pathology compared to the single deficiencies, indicating that the EGRESS complex works as a functional unit. Mutagenesis experiments showed that the second luminal loop of CLN6 is required for the interaction of CLN6 with the enzymes but dispensable for interaction with CLN8, and in vitro and in vivo studies showed that CLN6 deficiency results in inefficient ER export of lysosomal enzymes and diminished levels of the enzymes at the lysosome. These results identify CLN6 and the EGRESS complex as key players in lysosome biogenesis and shed light on the molecular etiology of Batten disease caused by defects in CLN6.

## INTRODUCTION

Lysosomes contain more than 50 soluble hydrolytic enzymes that mediate the degradation of macromolecules according to various catabolic programs. Lysosomal enzymes are synthesized at the endoplasmic reticulum (ER) and transferred to the endolysosomal system via the secretory route (1, 2). Whereas the post-Golgi trafficking of lysosomal enzymes has been amply characterized (3), the early, pre-Golgi stages of lysosomal enzyme trafficking are only partially understood. We have recently reported that ER-to-Golgi transfer of newly synthesized lysosomal enzymes is mediated by the cargo receptor CLN8 (ceroid lipofuscinosis, neuronal, 8) (4). CLN8 is a ubiquitously expressed, multi-pass membrane protein that forms homodimers and localizes in the ER and the ER-Golgi intermediate compartment (5, 6). CLN8 interacts with newly synthesized lysosomal enzymes in the ER, transfer them to the Golgi via COPII vesicles, and recycles back to the ER via COPI vesicles (4). Absence of CLN8 results in inefficient ER exit and decreased levels of lysosomal enzymes in mouse tissues and patient-derived cells (4), resulting in a subtype of Batten disease or neuronal ceroid lipofuscinosis (NCL) (7, 8).

NCLs are a heterogeneous group of 13 genetically distinct progressive encephalopathies mainly presenting during infancy and childhood. The overall incidence of NCLs in the US is estimated to be 1:12,500 live births (9). NCLs are classified as lysosomal storage disorders because a prominent feature emerging from the pathological analysis of patients’ tissues is the intralysosomal accumulation of ceroid lipopigment (7). A clear link between protein dysfunction and lysosomal pathology in NCLs is established by the observation that four NCL subtypes are caused by mutations in one of four lysosomal enzymes (PPT1, TPP1, CTSD, CTSF). In addition, at least four other NCL genes encode proteins that localize and presumably function in the lysosome (10). Impaired trafficking of lysosomal enzymes due to CLN8 deficiency is another example linking impaired protein function to NCL molecular pathology (4).

The ubiquitously expressed, multi-pass membrane protein CLN6 (ceroid lipofuscinosis, neuronal, 6; OMIM 601780) is the only other NCL protein that resides in the ER (11, 12). Remarkably, the clinical features of NCL patients with mutations in *CLN6* or *CLN8* are strikingly similar (13, 14). Commonly associated symptoms include cognitive decline, seizures, retinopathy and gait difficulties with patients first reporting to the clinic between two and eight years of age (14–16). Naturally occurring mouse models for CLN6 and CLN8 diseases have been widely characterized and have been shown to mimic many of the disease phenotypes (17). Like human patients, mutant mice die prematurely and exhibit dysfunctional lysosomal metabolism in multiple tissues and organs (18, 19). The underlying defective molecular pathway linking CLN6 deficiency to lysosomal dysfunction and NCL is unclear, as the molecular function of CLN6 is yet to be characterized (20). Recent reports show alterations in metal homeostasis pathways— mostly accumulation of zinc and manganese—and aberrant cell signaling related to Akt/GSK3 and ERK/MAPK pathways as characteristic features of CLN6 disease (21). Impaired stability and function of the CLN6 interactor, collapsin response mediator protein-2 (CRMP-2), has also been associated with altered neurite maturation in CLN6 disease, possibly contributing to neuronal dysfunction and encephalopathy (22). While these reports indicate associations between tissue pathology and disease symptoms, however, the function of CLN6 and its relationship with lysosomal pathways has remained elusive.

Based on the similarities between the clinical features associated with CLN6 and CLN8 deficiencies and the partially overlapping subcellular localization of the two proteins, we hypothesized that CLN6 also functions in ER-to-Golgi transfer of lysosomal enzymes. Here we investigated whether CLN6 acts as a cargo receptor for lysosomal enzymes and whether CLN6 function is redundant with, or dependent on, the function of CLN8. Our results show that CLN6 and CLN8 are obligate partners for the recruitment of newly synthesized lysosomal enzymes at the ER and that, differently from CLN8, CLN6 is not loaded into COPII vesicles but is retained in the ER, presumably to serve additional cycles of enzyme recruitment. We determine that the second luminal loop of CLN6 is required for the interaction of CLN6 with the enzymes and that CLN6 deficiency results in inefficient ER export of lysosomal enzymes and diminished levels of the enzymes at the lysosome. These results identify CLN6 as a key protein implicated in the biogenesis of lysosomes and shed light on the molecular etiology of Batten disease caused by defects in CLN6.

## RESULTS

### Loss of CLN6 does not aggravate pathology of CLN8-deficient mice

To determine whether CLN6 and CLN8 have redundant functions we first conducted studies by using the *Cln6^nclf^* and *Cln8^mnd^* mouse lines (henceforth referred to as *Cln6^−/−^* and *Cln8^−/−^*), which carry early frameshift mutations in *Cln6* and *Cln8* genes, respectively, resulting in the complete absence of CLN6 and CLN8 proteins (11, 19, 23, 24). Life span analysis showed that *Cln6^−/−^* and *Cln8^−/−^* mice had a slightly different survival rate: The median survival of *Cln6^−/−^* mice was 450 days, while the median survival of *Cln8^−/−^* mice was 302 days (log-rank test; z = 7.07, *P* < 0.001), in line with previous reports (18, 19, 23). Double KO mice had a median survival of 290 days, which was indistinguishable from that of *Cln8^−/−^* mice (log-rank test; z = 2.24, p = 0.025) (Figure 1A). Thus, simultaneous deficiency of CLN6 and CLN8 does not accelerate disease progression compared to CLN8 single deficiency.

**Figure 1.**
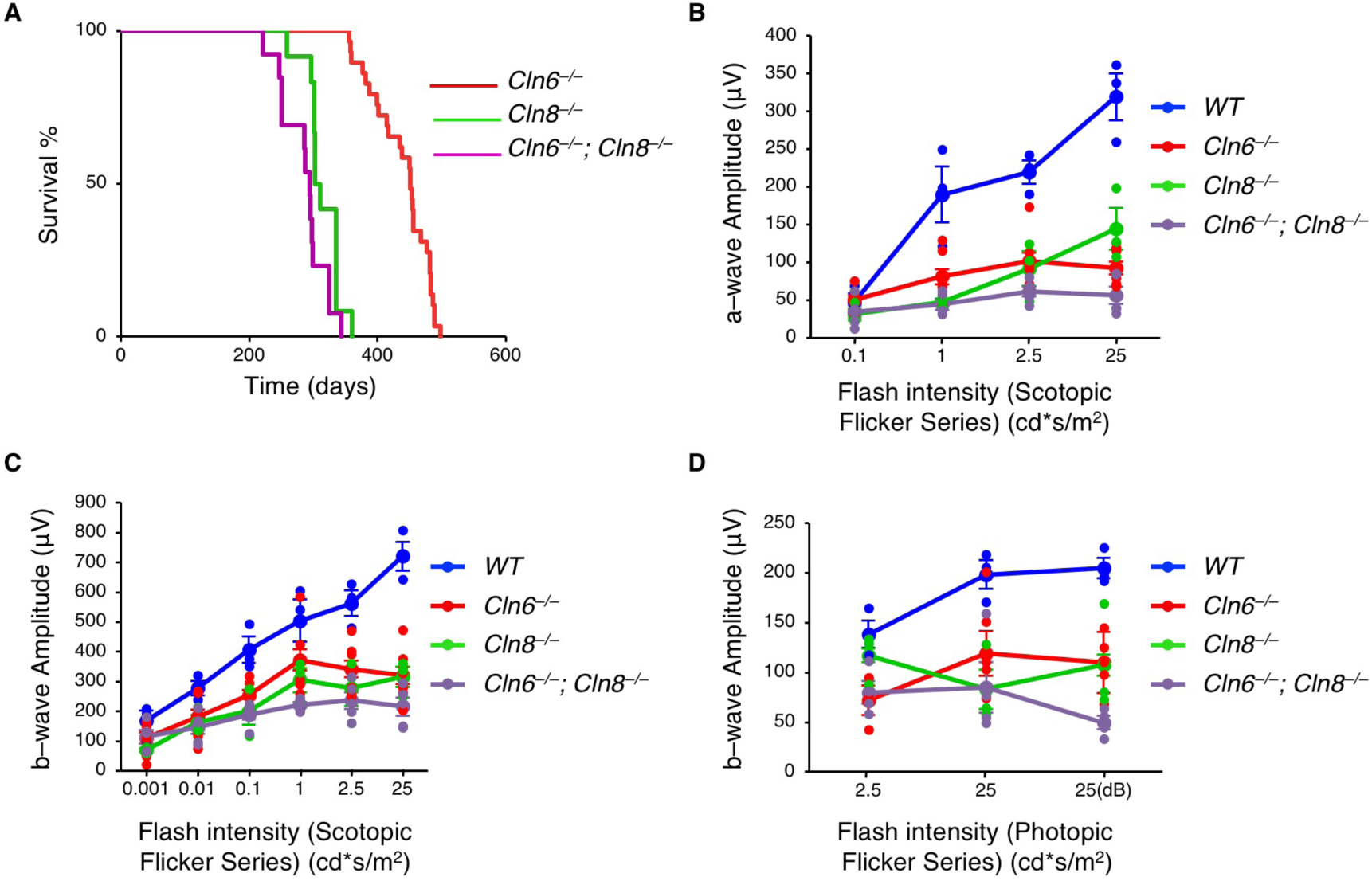
Combined deficiency of CLN6 and CLN8 in mice does not accelerate disease progression compared to single deficiencies. **(A)** Life span analysis of *Cln6*^−/−^ (red), *Cln8*^−/−^ (green) and *Cln6*^−/−^; *Cln8*^−/−^ (magenta) mice. *n* = 16-26 mice/genotype. **(B)** Graph of a-wave amplitudes of scotopic ERG waveforms from WT (blue), *Cln6*^−/−^ (red), *Cln8*^−/−^ (green), and *Cln6*^−/−^; *Cln8*^−/−^ (magenta) mice in response to a series of light stimuli. **(C)** Graph of b-wave amplitudes of scotopic ERG waveforms from WT (blue), *Cln6*^−/−^ (red), *Cln8*^−/−^ (green), and *Cln6*^−/−^; *Cln8*^−/−^ (purple) mice. **(D)** Graph of b-wave amplitudes of photopic ERG waveforms from WT (blue), *Cln6*^−/−^ (red), *Cln8*^−/−^ (green), and *Cln6*^−/−^; *Cln8*^−/−^ (purple) mice. Error bars in (B-C) are SEM.

A notable phenotype described in most mouse models of Batten disease is retinal degeneration, which eventually leads to complete blindness (25, 26). Both *Cln6^−/−^* and *Cln8^−/−^* mouse lines display early-onset loss of vision that can be quantified by electroretinogram (ERG) analysis (25, 27). To compare the visual function of single and double KO mouse lines, we performed scotopic and photopic ERGs to assess the function of rods and cones, respectively. Scotopic ERG was performed by measuring a-wave (photoreceptor) and b-wave (inner retina) amplitudes. The results from both scotopic (Figure 1, B and C) and photopic (Figure 1D) ERGs showed that, while all mutants displayed significant differences compared to age-matched wild-type (WT) mice, there was no difference in the comparison of the single and double KO lines. Together, life span and ERG analyses suggest that CLN6 and CLN8 do not have redundant functions. We therefore set up to investigate whether they work in the same biological process— as partners of a protein complex or in sequential steps of a common pathway.

### CLN6 deficiency results in the depletion of various lysosomal enzymes from the lysosomal compartment

We recently demonstrated that, in absence of CLN8, lysosomes have diminished levels of various soluble lysosomal enzymes (4). To investigate whether CLN6 deficiency also results in defective lysosomal composition, we isolated lysosome-enriched fractions from the livers of pre-symptomatic, 6-week-old *Cln6^−/−^* mice and age-matched WT mice using a discontinuous Nycodenz-sucrose gradient as described (28). Immunoblot analysis for the lysosomal membrane protein LAMP1 confirmed lysosomal enrichment in the collected fractions and showed no obvious changes in LAMP1 signal between WT and *Cln6^−/−^* samples (Supplemental Figure 1A). We confirmed lysosomal enrichment by performing enzyme assay for β-hexosaminidase, a lysosomal enzyme that is not affected by CLN8 deficiency (4) and that also did not show changes upon deficiency of CLN6 (Supplemental Figure 1B). We then performed immunoblot analysis for a set of enzymes for which antibodies able to recognize the mouse proteins are available. The results showed a general reduction in enzyme levels in the lysosomal fraction from *Cln6^−/−^* mice (Figure 2, A and B). Consistent with these results, we observed a decrease in lysosomal enzyme activities using proteins extracted from the lysosome-enriched fraction of *Cln6^−/−^* mice (Figure 2C). Real-time qPCR using liver RNAs, however, showed slightly increased transcription of lysosomal enzymes in *Cln6^−/−^* mice compared to WT controls (Figure 2D). In addition, confocal microscopy analysis of mouse embryonic fibroblasts (MEFs) from WT and *Cln6^−/−^* mice showed decreased overlap of CTSD and PPT1 with the lysosomal marker LAMP1 in absence of CLN6, whereas no obvious changes in the overlap of these enzymes with the ER protein KDEL were found (Supplemental Figure 1C-F). Together, these results show an absence of major defects in the expression of lysosomal enzymes and indicate that the cause of the observed lysosomal enzyme depletion must be post-translational.

**Figure 2.**
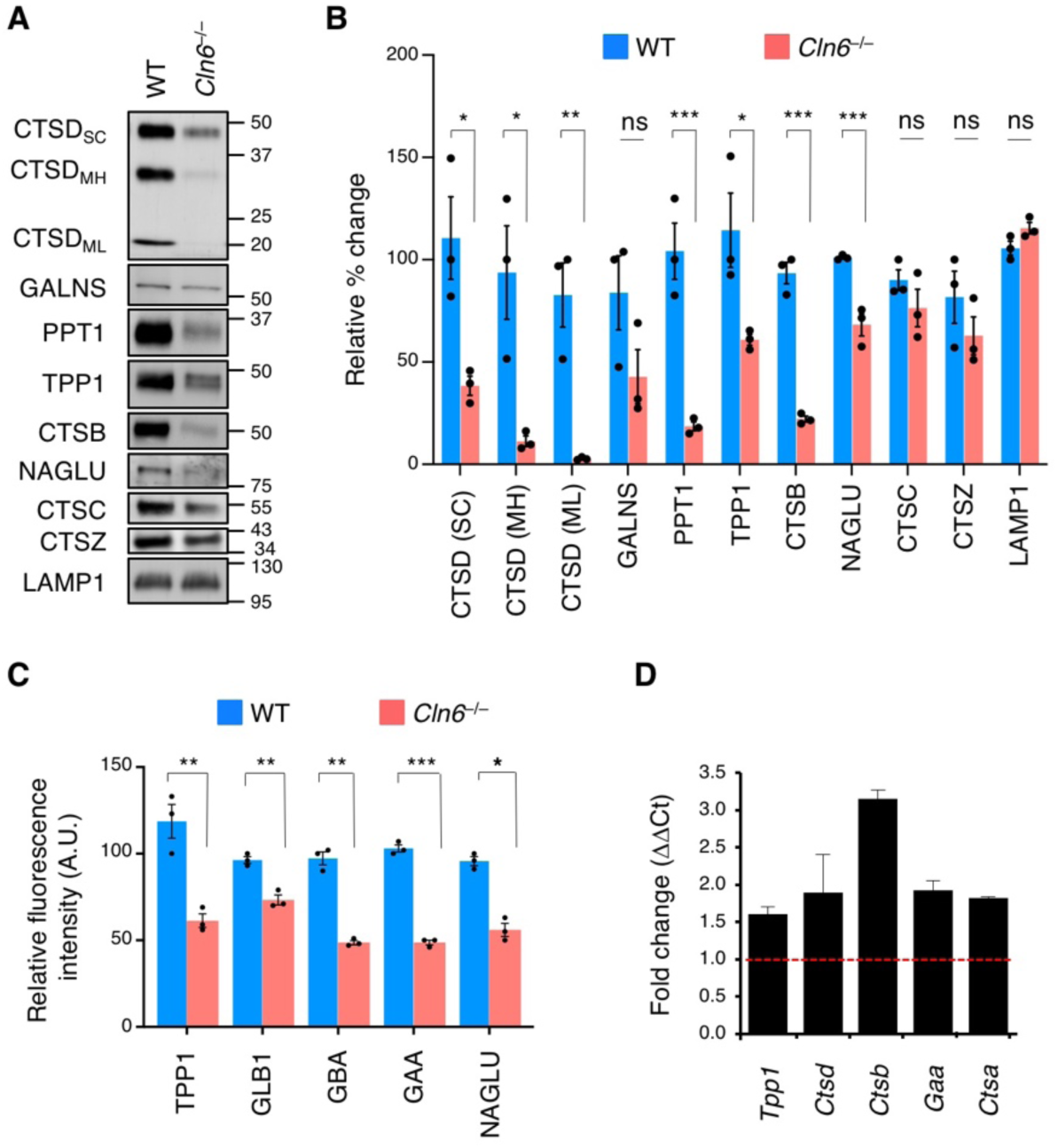
CLN6 deficiency results in the depletion of various lysosomal enzymes from the lysosomal compartment. **(A)** Immunoblot analysis of lysosome-enriched fractions confirming depletion of lysosomal enzymes in *Cln6*^−/−^ mice compared to WT mice. CTSD_SC_, single chain processed form; CTSD_MH_, mature heavy form; CTSD_ML_, mature light form. **(B)** Band intensities were quantified and normalized to LAMP1. Data are means ± SEM (*n* = 3, *P < 0.05, **P < 0.01). **(C)** Enzymatic assay of TPP1, GBA, GLB1, GAA and NAGLU in lysosome-enriched fractions from WT and *Cln6*^−/−^ mice. Activity is expressed as relative fluorescence units compared to WT samples. Data are means ± SEM (*n* = 3, *P < 0.05, **P < 0.01). **(D)** Expression analysis of lysosomal genes in the liver of 6-weeks-old WT and *Cln6*^−/−^ mice. Shown are expression levels of genes in *Cln6*^−/−^ mice expressed as fold change of levels in WT mice, normalized to the housekeeping gene *Cyclophilin*. Data are means ± SEM (*n = 3*).

### CLN6 interacts with CLN8 but does not traffic to the Golgi complex

We next investigated whether CLN6 interacts with CLN8 by using bimolecular fluorescence complementation (BiFC), an assay in which proteins are tagged with either the N- or the C-terminus of YFP (Y1 and Y2, respectively), and fluorescence is emitted by reconstituted YFP upon interaction of the two tagged proteins (29). BiFC assays performed by co-transfecting either CLN6-Y1 with Y2-CLN8 or CLN6-Y2 with Y1-CLN8 showed interaction between CLN6 and CLN8 under either Y1/Y2 configuration in HeLa cells (Figure 3A, Supplemental Figure 2A) and mouse embryonic fibroblasts (Supplemental Figure 2B). As a control, we verified that CLN6 does not interact with lipase maturation factor 1 (LMF1) (Figure 3A), an ER transmembrane protein that acts as a cargo receptor for lipoprotein lipase (30). Co-staining with KDEL showed colocalization with the reconstituted fluorescence, indicating that CLN6 and CLN8 interact at the ER (Figure 3B; Pearson correlation coefficient = 0.76 ± 0.08, *n* = 10 cells). Co-IP experiments using tagged proteins expressed in HEK293-T cells confirmed that CLN6 interacts with CLN8 (Supplemental Figure 2C) but not with LMF1 (Supplemental Figure 2D). In addition, we generated a cell line in which endogenous CLN8 is fused to a myc tag by using CRISPR-Cas9 genome editing. Co-IP experiments confirmed interaction of CLN6 with endogenous CLN8 (Figure 3C).

**Figure 3.**
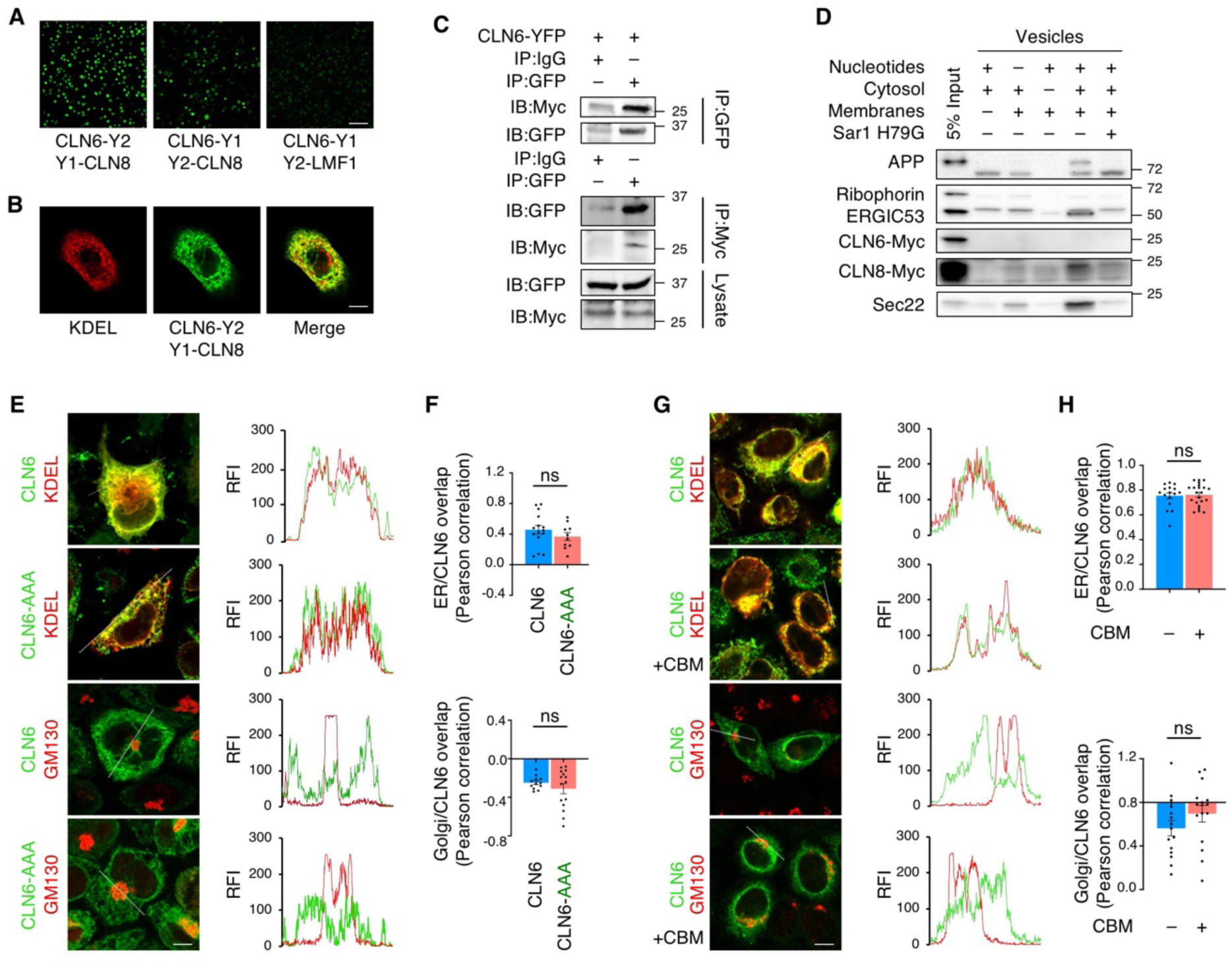
CLN6 interacts with CLN8 in the ER and does not traffic to the Golgi complex. **(A)** Shown is a live BiFC assay of CLN6 with CLN8 in the indicated YFP configurations. Green signals (reconstituted YFP) represent CLN6-CLN8 interaction. LMF1 is used as a negative control for interaction with CLN6. Scale bar: 200 µm. **(B)** Confocal microscopy analysis showing colocalization of reconstituted CLN6-Y2/Y1-CLN8 BiFC signal with the ER marker KDEL. Scale bar: 20 µm. **(C)** Co-IP analysis of transiently expressed, Y2-tagged CLN6 and endogenous, myc-tagged CLN8. The lysates were immunoprecipitated with both myc and GFP antibodies in separate experiments and analyzed by immunoblotting with the indicated antibodies. IgG antibodies were used as a control. Input represents 10% of the total cell extract used for IP. (**D**) In vitro COPII vesicle budding assay on digitonin-treated HeLa cell membranes incubated with the indicated combinations of ATP regenerating system, rat liver cytosol, collected donor membranes, and dominant-negative Sar1H79G; 5% input of donor membranes is included. (**E**) Confocal microscopy analysis showing that CLN6 resides in the ER upon mutagenesis of a potential retrieval/retention signal (RRR to AAA). Trace outline is used for line-scan analysis of Relative Fluorescence Intensity (RFI) of CLN6, GM130, and KDEL signals. Scale bar: 10 µm. (**F**) Pearson correlation analysis of the co-localization extent of full-length CLN6 or CLN6-AAA with KDEL or GM130. *n* > 20 images/experiment. (**G**) Confocal microscopy showing that CLN6 resides in the ER upon treatment with CBM. Trace outline is used for RFI line-scan analysis of CLN6, GM130, and KDEL signals. Scale bar: 20 µm. (**H**) Pearson correlation analysis of the co-localization extent of CLN6 with KDEL or GM130 upon treatment with CBM. *n* >20 images/experiment.

Additional confocal microscopy analyses showed that both CLN6 and CLN8 colocalize with Sec16L, a marker for ER exit sites (Supplemental Figure 2E). To test whether CLN6 is loaded into COPII vesicles we performed an *in vitro* COPII vesicle budding assay. We incubated membranes from myc-CLN6-transfected HeLa cells with rat liver cytosol and nucleotides for 1 hr at 30°C. Newly formed vesicles were collected and blotted for CLN6 and CLN8. Three known COPII cargo proteins—APP, ERGIC-53 and Sec22—were used as positive controls (31, 32), and Ribophorin I, an ER-resident protein excluded from COPII vesicles (33), was used as a negative control. Unlike CLN8, APP, and ERGIC-53, which were detected in COPII vesicles, CLN6 was excluded from COPII vesicles, similar to Ribophorin I (Figure 3D). Addition of Sar1 H79G, a dominant negative mutant that inhibits COPII vesicle formation (31), blocked the inclusion of APP, Sec22, ERGIC-53, and CLN8 into newly formed vesicles, demonstrating that the process is specifically dependent on COPII.

In agreement with CLN6 exclusion from COPII vesicles, mutation of a putative ER retention/retrieval signal (^5^RRR mutated to AAA) (34) present in the cytosol-facing N-terminus of CLN6 did not change the subcellular localization of CLN6 (Figure 3, E and F), consistent with previous observations (20). Because other unknown protein signals could mediate retrieval of CLN6 from the Golgi to the ER, we also inhibited Golgi-to-ER retrograde trafficking using 1,3-cyclohexanebis(methylamine) (CBM), a chemical inhibitor of COPI-mediated vesicular transport (35). CBM treatment resulted in a change of CLN8 localization to the Golgi (Supplemental Figure 2, F and G) as previously observed (4), whereas it did not result in any discernible overlap of CLN6 with the Golgi marker, confirming exclusive localization of CLN6 in the ER (Figure 3, H and I). These results are in agreement with prior work indicating that CLN6 does not localize to the Golgi (20).

### CLN8 trafficking to the Golgi is uncoupled from CLN6-CLN8 interaction

We next investigated whether the subcellular localizations of CLN6 and CLN8 are mutually dependent. We first tested whether abolishing Golgi-to-ER retrieval of CLN8 affects the subcellular localization of CLN6. To this end, we co-transfected myc-tagged CLN6 with either full-length Y2-CLN8 or the retrieval-deficient CLN8 mutant Y2-CLN8dK (4). Confocal microscopy showed that, in either case, CLN6 localized at the ER in HEK293-T cells (Figure 4, A and B). To avoid interaction between CLN6 and endogenous CLN8, we repeated the test in a cell line in which CLN8 was knocked-out (*CLN8*^−/−^) (4) and obtained a similar result (Supplemental Figure 2, H and I). We then tested whether CLN8 can traffic to the Golgi independently of CLN6. To this aim, we generated *CLN6^−/−^* cells by CRISPR-Cas9 genome editing (Supplemental Figure 3, A-C); as a control, we confirmed depletion of lysosomal enzymes in these cells, which could be rescued by *CLN6* re-introduction (Supplemental Figure 3D). Confocal microscopy showed that retrieval-deficient Y2-CLN8dK localizes to the Golgi in *CLN6^−/−^* cells (Figure 4, C and D), indicating that CLN8 trafficking is uncoupled from interaction with CLN6.

**Figure 4.**
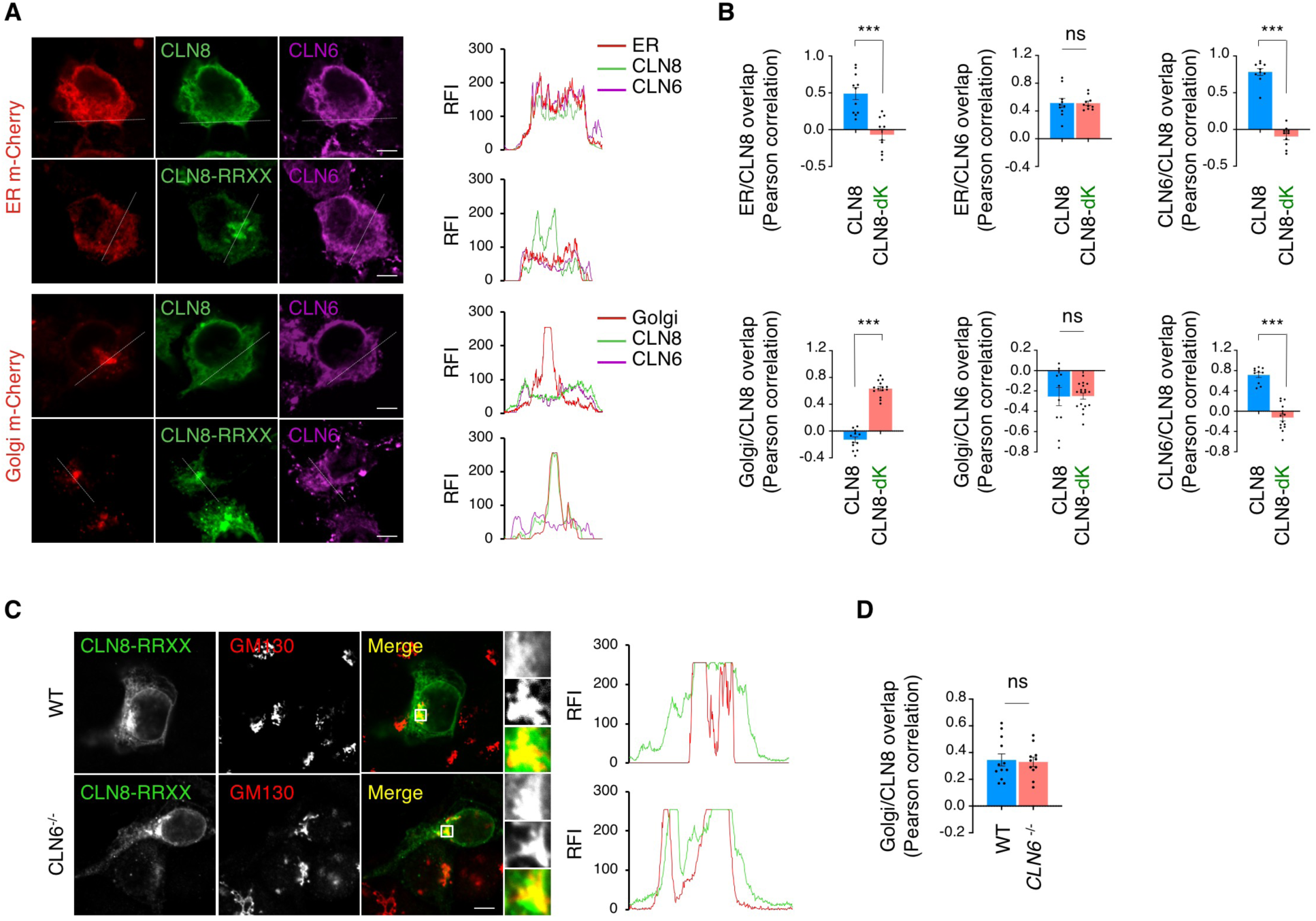
The subcellular localizations of CLN6 and CLN8 are uncoupled from CLN6-CLN8 interaction. (**A**) Confocal microscopy analysis showing ER localization of CLN6 and Golgi localization of CLN8 upon co-transfection of full-length CLN6 and retrieval-deficient CLN8. Trace outline is used for RFI line-scan analysis of CLN8, CLN6, Golgi-cherry and ER-cherry signals. Scale bar: 10 µm. *n* >20 images/experiment. (**B**) Pearson correlation analysis of the co-localization extent of CLN6 and CLN8 constructs with KDEL and GM130. (**C**) Confocal microscopy showing that retrieval-deficient CLN8 (CLN8-RRXX, green signal) has partial colocalization with the Golgi marker GM130 (red) both in WT and *CLN6*^−/−^ cells. Scale bar: 20 µm. Inset magnifications (5x) are reported. (**D**) Pearson correlation analysis showing partial co-localization of retrieval deficient (CLN8-RRXX, green signal) with the Golgi marker GM130 in WT and *CLN6*^−/−^ cells. *n* >20 images/experiment.

### CLN6 interacts with lysosomal enzymes and the interaction requires CLN6’s second luminal loop

Next, we examined whether CLN6 interacts with the lysosomal enzymes. Co-immunoprecipitation assay using myc-tagged CLN6 and a set of YFP-tagged enzymes showed that CLN6 interacts with the precursor forms of enzymes that we previously characterized as CLN8 interactors (CTSD, PPT1, TPP1, and GALNS), while did not show interaction with HEXB, which also did not interact with CLN8 (4) (Figure 5A). Co-IP of CLN6 with the non-lysosomal secretory proteins AGN and TGF-1β resulted in the absence of any detectable interaction, thus supporting the notion that CLN6 interactions are specific. To gain insight into the domains of CLN6 involved in the interaction with the enzymes, we analyzed the structure and conservation of the CLN6 protein. CLN6 has a cytosolic N-terminus, seven transmembrane domains, and a C-terminus in the ER lumen (16, 20, 36). The cytosolic and luminal loops connecting the transmembrane domains are all small (<15 amino acids), with the exception of the second luminal loop that is 48 amino-acid long (Figure 5B). Evolutionary constrained region analysis (37) using 26 vertebrate species showed that CLN6’s second luminal loop (comprised between transmembrane domains 3 and 4) is the most constrained region of the protein (Supplemental Figure 4). We therefore hypothesized that this loop is involved in the interaction of CLN6 with the lysosomal enzymes. To test this hypothesis, we generated a CLN6 construct lacking the second luminal loop (CLN6ΔL2) by deleting the amino acids from position 135 to position 175 (Supplemental Figure 5, A and B). We first verified that the CLN6ΔL2 protein, like full-length CLN6, localizes at the ER (Supplemental Figure 5C). Given that CLN6 forms homodimers (Figure 5C) (20), we also tested whether CLN6ΔL2 is able to dimerize with full-length and ΔL2 CLN6 proteins. BiFC assays showed that full-length, YFP-tagged CLN6 forms a dimer (CLN6-Y1/CLN6-Y2) that is detectable by BiFC coupled with either confocal microscopy (Figure 5D) or flow cytometry (Supplemental Figure 5, D and E). The CLN6-Y1/CLN6-Y2 dimer correctly localized in the ER as detected by confocal microscopy (Figure 5E). BiFC analysis showed that CLN6ΔL2 retains the ability to form dimers with full-length CLN6 (CLN6ΔL2-Y1/CLN6-Y2 and CLN6ΔL2-Y2/CLN6-Y1) as well as with itself (CLN6ΔL2-Y1/CLN6ΔL2-Y2) (Figure 5F). Thus, the second luminal loop of CLN6 is dispensable for protein stability and self-interaction. We then performed co-IP assay to test whether CLN6ΔL2 is able to interact with the lysosomal enzymes. Pull-down of myc-tagged CLN6ΔL2 followed by immunoblotting for Y1-tagged lysosomal enzymes (detectable with an anti-GFP antibody) showed that deletion of the second loop of CLN6 disrupts the interaction with the tested enzymes (Figure 5G). Thus, the second loop of CLN6 is required for the interaction of CLN6 with the lysosomal enzymes.

**Figure 5.**
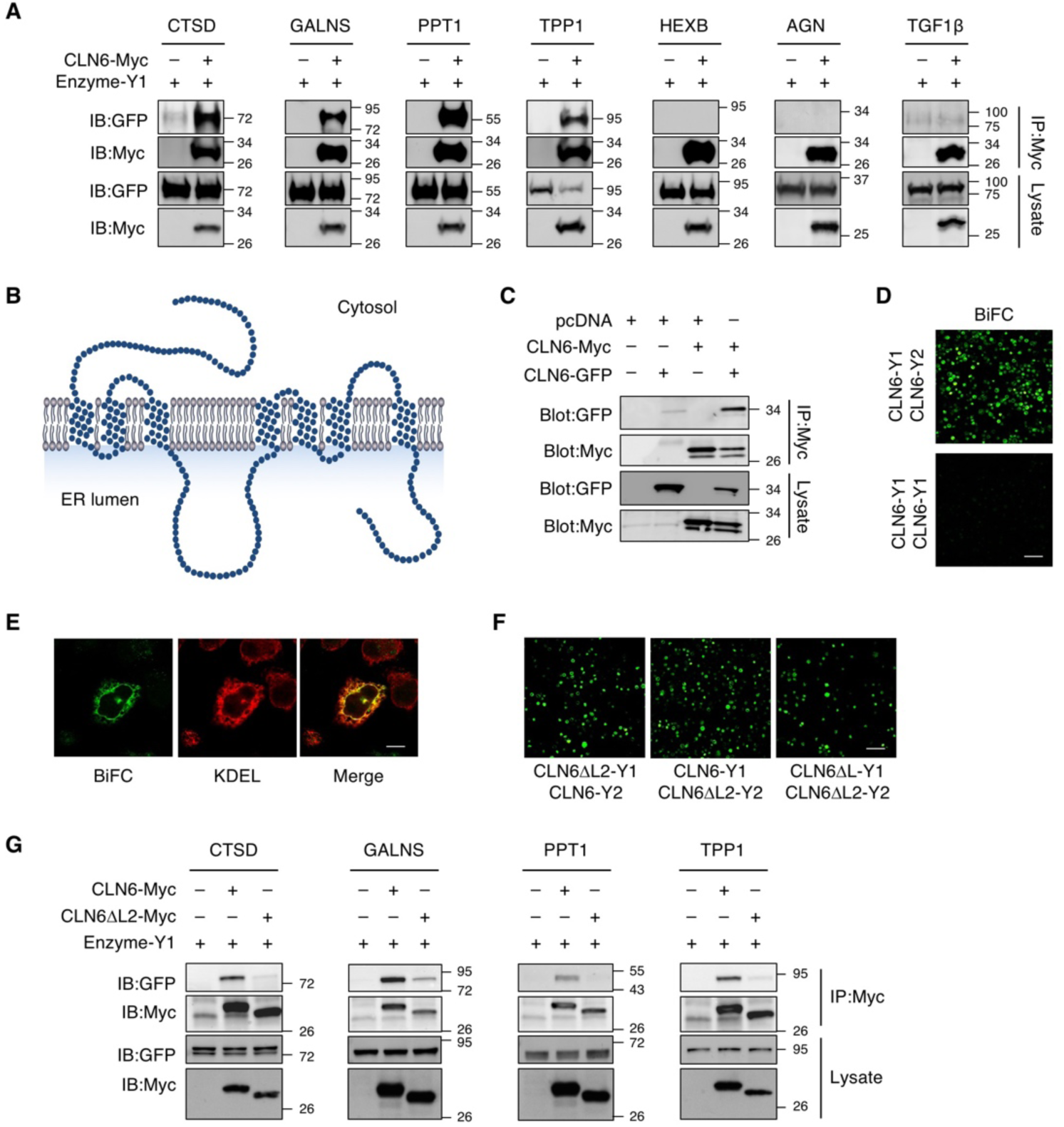
The second luminal loop of CLN6 is necessary for the interaction of CLN6 with the lysosomal enzymes. **(A)** Co-IP analysis of CLN6 and lysosomal enzymes. Proteins were transiently expressed in HEK293-T cells, and immunoprecipitates were analyzed by immunoblotting with the indicated antibodies. Molecular marker analysis indicates that CLN6 interacts with the enzymes’ precursors. Input represents 10% of the total cell extract used for IP. **(B)** Schematic representation of CLN6 protein. **(C)** Co-IP analysis of Y2-tagged CLN6 and myc-tagged CLN6. The proteins were transiently expressed in HEK293-T cells, and immunoprecipitates were analyzed by immunoblotting with the indicated antibodies. Input represents 10% of the total cell extract used for IP. **(D)** Shown is a live-BiFC assay of CLN6-Y1 with CLN6-Y2 in HeLa cells; expression of CLN6-Y1 is used as a negative control. Scale bar: 200 µm. **(E)** Confocal microscopy showing colocalization between reconstituted BiFC signal from CLN6-Y1/CLN6-Y2 dimerization (green) and the ER marker KDEL (red). Scale bar: 20 µm. **(F)** Shown are live-BiFC assays of CLN6ΔL2-Y1 with CLN6-Y2, CLN6-Y1 with CLN6ΔL2-Y2, and CLN6ΔL2-Y1 with CLN6ΔL2-Y2 in HeLa cells. Scale bar: 200 µm. **(G)** Co-IP analysis of CLN6ΔL2 and lysosomal enzymes. Proteins were transiently expressed in HEK293-T cells, and immunoprecipitates were analyzed by immunoblotting with the indicated antibodies. Input represents 10% of the total cell extract used for IP.

### CLN6 and CLN8 are mutually necessary for their interaction with lysosomal enzymes

Next, we used *CLN6^−/−^* and *CLN8^−/−^* cells to investigate whether CLN6 and CLN8 are mutually necessary for their interaction with lysosomal enzymes. Co-IP assays showed that CLN6 is unable to interact with the lysosomal enzymes in *CLN8^−/−^* cells (Figure 6A); similarly, the interaction of CLN8 with the lysosomal enzymes is abolished in *CLN6^−/−^* cells (Figure 6B). In the absence of CLN6, however, the ability of CLN8 to form homodimers was not abolished as seen by BiFC analysis followed by flow-cytometry (Figure 6C). Similarly, CLN6 was able to form homodimers in *CLN8^−/−^* cells (Figure 6D).

**Figure 6.**
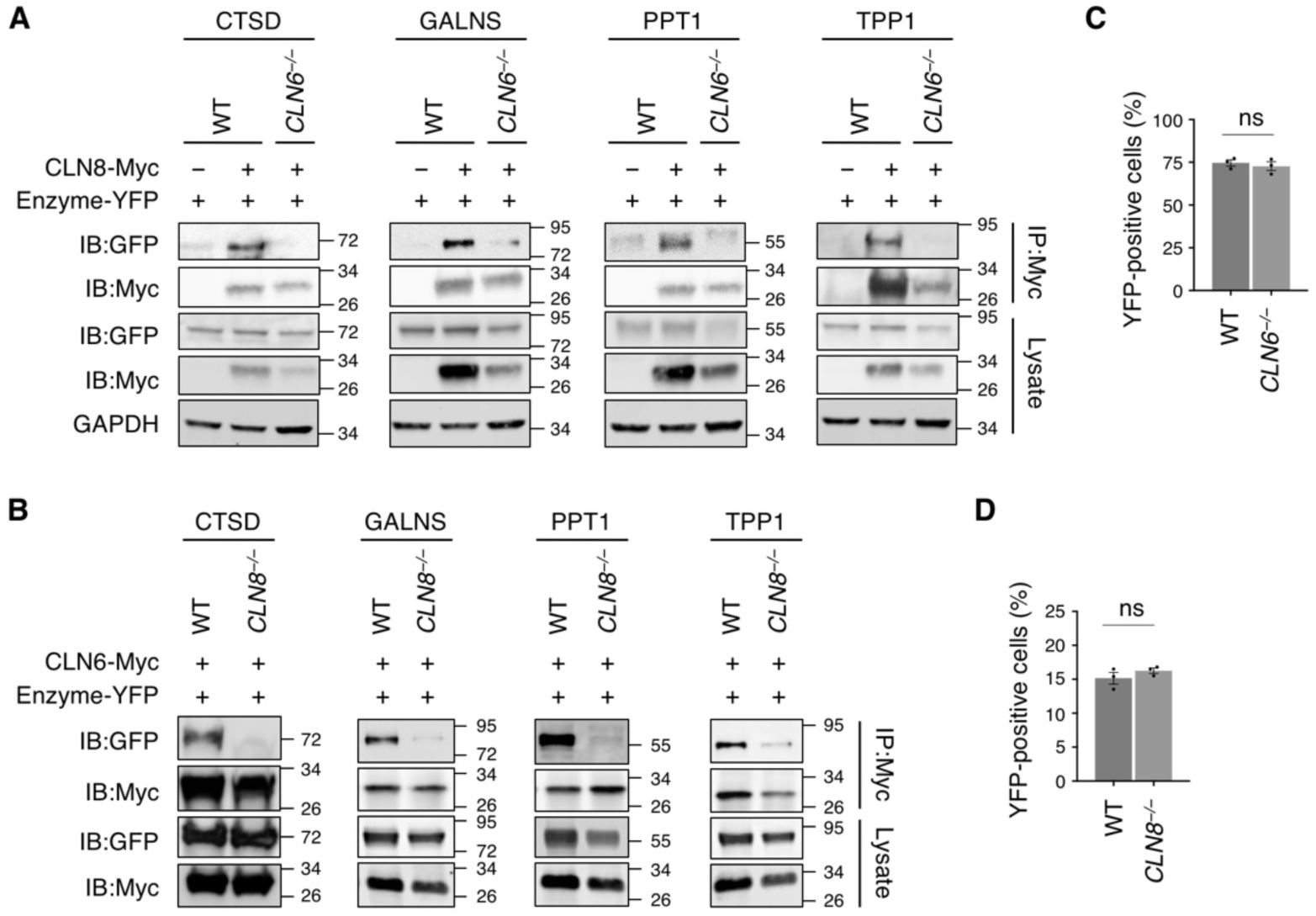
CLN6 and CLN8 are mutually necessary for their interaction with lysosomal enzymes. (**A**) Co-IP analysis of myc-tagged CLN8 and Y2-tagged lysosomal enzymes (TPP1, CTSD, PPT1, and GALNS). Vectors were transiently transfected in WT and *CLN6^−/−^* HEK293-T cells, and immunoprecipitates were analyzed by immunoblotting with the indicated antibodies. Input represents 10% of the total cell extract used for IP. **(B)** Co-IP analysis of myc-tagged CLN6 and Y2 tagged lysosomal enzymes (TPP1, CTSD, PPT1, and GALNS). Vectors were transiently transfected in WT and *CLN8^−/−^* HEK293-T cells, and immunoprecipitates were analyzed by immunoblotting with the indicated antibodies. Input represents 10% of the total cell extract used for IP. (**C**) Flow cytometry quantification of Y1-CLN8/Y2-CLN8 BiFC signal in WT and *CLN6*^−/−^ cells. **(D)** Flow cytometry quantification of CLN6-Y1/CLN6-Y2 BiFC signal in WT and *CLN8*^−/−^ cells.

Based on these results, we conclude that CLN6 and CLN8 are obligate partners in the recruitment of newly synthesized lysosomal enzymes in the ER, and that the subsequent transfer of enzymes to the Golgi is mediated by CLN8 only. Although CLN6 is not loaded in COPII vesicles along with CLN8 and lysosomal enzymes, the observed decreased levels of enzymes in the lysosomal compartment upon CLN6 deficiency and the fact that CLN6 is essential for their recruitment in complex with CLN8 predict that, in absence of CLN6, ER-to-Golgi transfer of the enzymes may be inefficient. To test this hypothesis, we set out to monitor ER-to-Golgi trafficking of CTSD, PPT1 and GALNS by using the RUSH (Retention Using Selective Hooks) system (38) based on a recent report that showed that ER-to-Golgi transfer of CTSD occurs in a short time (< 30 min) and therefore requires a highly synchronized system for careful evaluation (39). In the RUSH system, the test protein is fused with a KDEL-tagged streptavidin binding protein (SBP), which enables retention of the test protein in the subcellular compartment of choice by simultaneous expression of an organelle-targeted hook protein that contains a streptavidin domain. The interaction between the test protein and the hook protein is stable and can be reversed by the addition of biotin to outcompete SBP binding. Biotin supplementation thus results in a synchronous release of the test protein, which can be monitored for relocation by time-course confocal microscopy. For our experiments, we co-expressed an ER-targeted hook protein (38) with SBP- and GFP-fused CTSD, PPT1, and GALNS. We used SBP-, GFP-fused lysozyme C (LyzC) (39) as a control protein. We monitored the subcellular localization of CTSD, PPT1, GALNS, and LyzC at 0, 5, 10, and 20 minutes from their biotin-induced synchronous release in WT and *CLN6^−/−^* cells. As a quantitative measure, we assessed the overlap of the enzyme signal with that of the Golgi marker GM130 at each time point. The results showed that the absence of CLN6 caused a significant delay in ER-to-Golgi trafficking of the tested lysosomal enzymes, but not of LyzC (Figure 7), thus confirming that the lack of a functional CLN6-CLN8 complex for the recruitment of lysosomal enzymes results in their inefficient exit from the ER.

**Figure 7.**
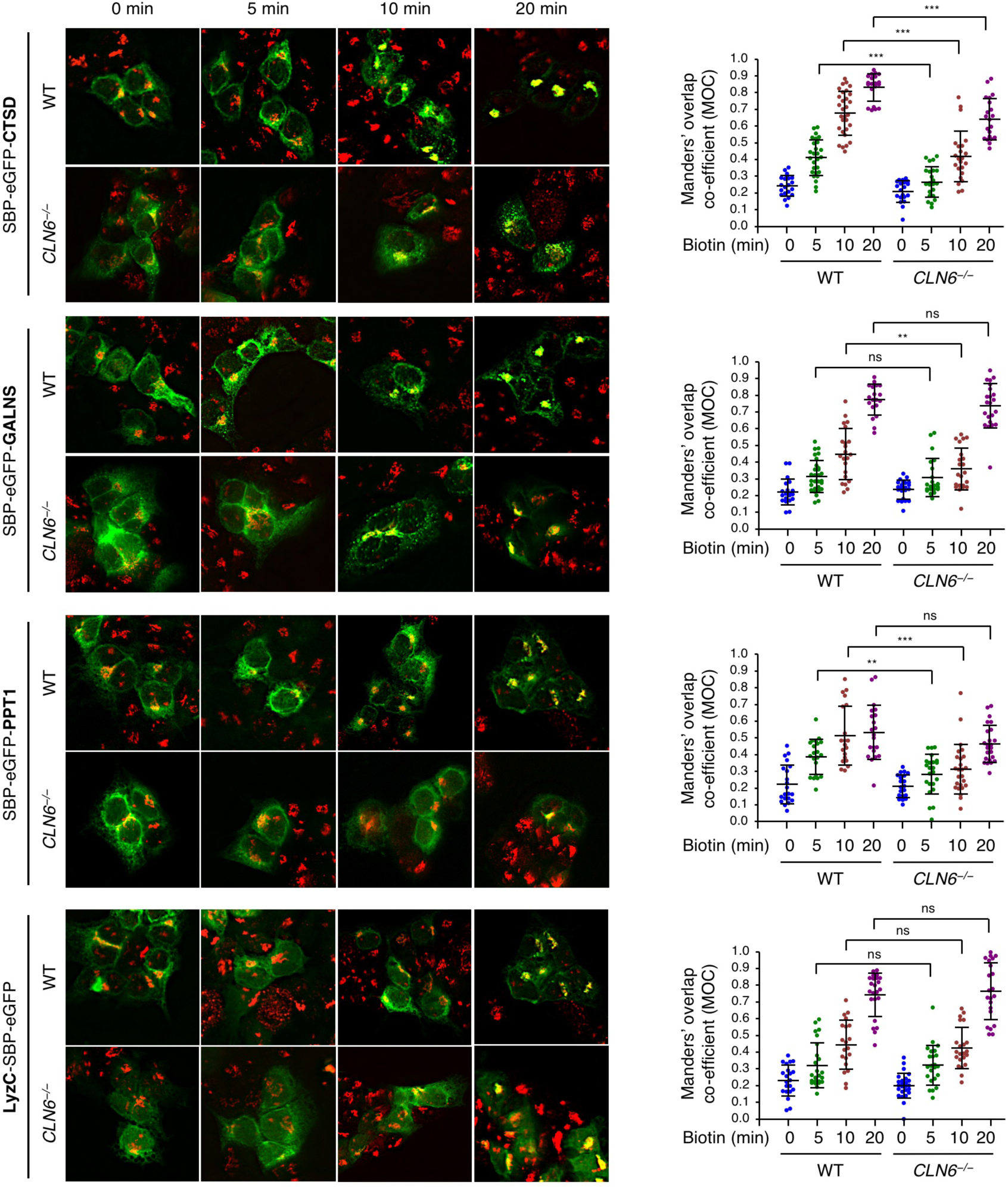
CLN6 deficiency impairs trafficking of lysosomal enzymes. Confocal microscopy analysis of WT and *CLN6^−/−^* HEK293-T cells transfected with plasmids expressing enzymes fused with streptavidin binding protein (SBP)-eGFP (SBP-eGFP-CTSD, SBP-eGFP-GALNS, SBP-eGFP-PPT1) and streptavidin-KDEL ‘‘anchor’’ that retains SBP-containing proteins in the ER. Shown are representative images of cells without addition of biotin (0 min) and at 5, 10, and 20 min from the addition of biotin. Lysozyme C (LysC-SBP-eGFP), a secretory protein that does not reside in the lysosome, is used as a negative control. Manders’ overlap coefficients measuring the degree of colocalization between the test protein (green signal) and the Golgi marker GM130 (red signal) are reported. Data are means ± SD (*n* > 20; *P < 0.05, **P < 0.01, ***P < 0.001; ns, not significant). Scale bars: 100 µm.

## DISCUSSION

This study identifies CLN6 as a key factor for the biogenesis of lysosomes by uncovering its function as an obligate component of a CLN6-CLN8 complex that recruits lysosomal enzymes at the ER to promote their Golgi transfer. We refer to this complex as EGRESS (<u>E</u>R-to-<u>G</u>olgi relaying of <u>e</u>nzymes of the ly<u>s</u>osomal <u>s</u>ystem) because of its role in recruiting lysosomal enzymes for Golgi delivery. The emerging model identifies a two-step process for the engagement of lysosomal enzymes in the ER and their transfer to the Golgi based on the cooperation between CLN6 and CLN8. In the first step, CLN6 and CLN8 form the EGRESS complex, which recruits the enzymes via an interaction that depends on both the large luminal loop of CLN6 (which is dispensable for the interaction with CLN8) and the large luminal loop of CLN8 (4). The absence of either protein makes the other unable to interact with the enzymes, indicating that the EGRESS complex is the minimal unit required for such interaction. In the second step, CLN8 is loaded into COPII vesicles to escort the enzymes to the *cis*-Golgi. It is possible that, during the loading into COPII vesicles, CLN6 is displaced by the interaction between CLN8 and COPII components, thus ending the loading cycle of the EGRESS complex. Upon Golgi transfer, the enzymes are subsequently trafficked to the lysosomes via selective transport from the *trans*-Golgi network, whereas the empty receptor is recycled back to the ER. If either component of the EGRESS complex is absent (or functionally defective), ER exit of the enzymes is inefficient, thereby resulting in enzyme depletion at the lysosome. A graphical depiction of inefficient enzyme exit from the ER upon CLN6 deficiency is reported in Figure 8.

**Figure 8.**
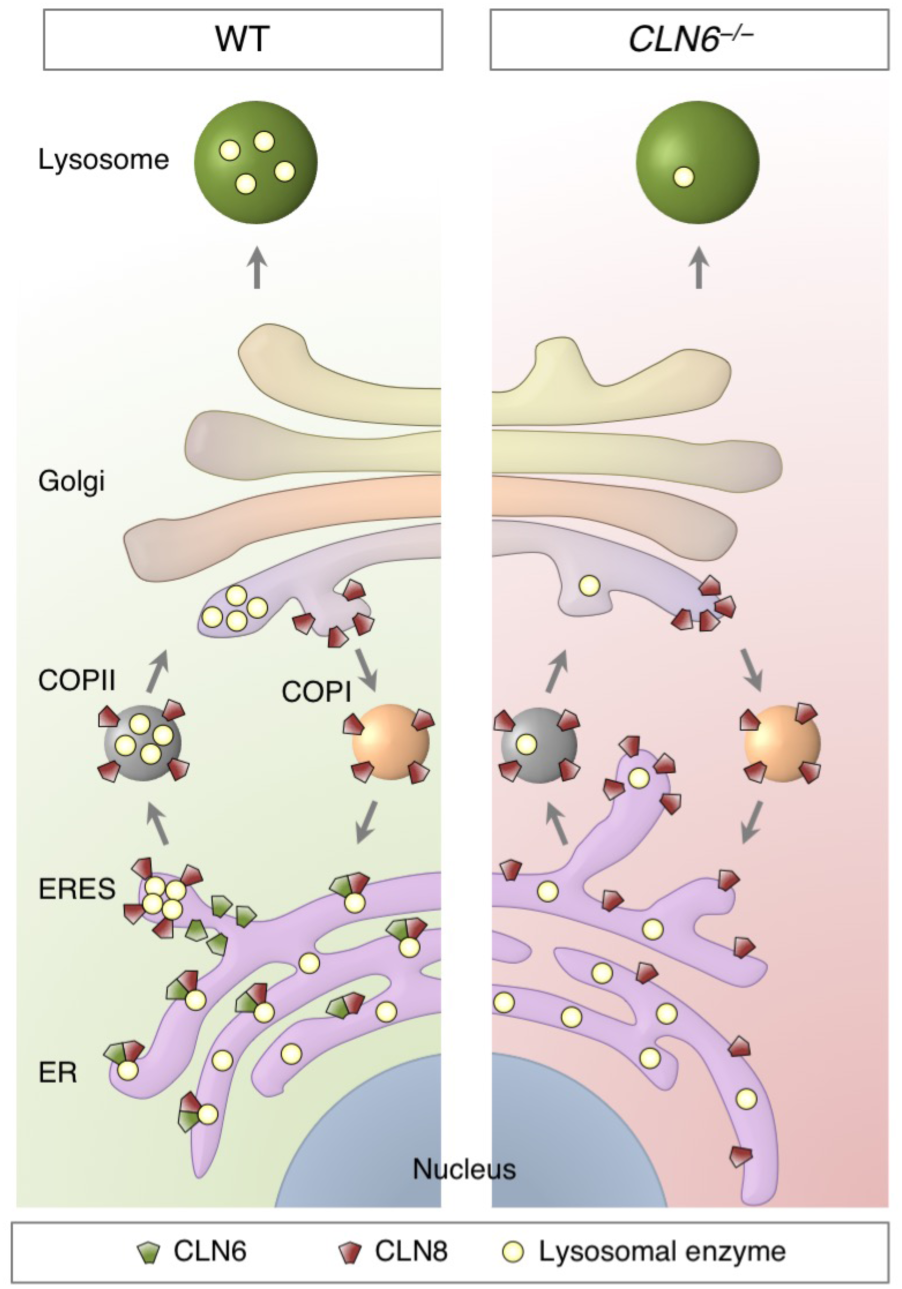
Schematic model of ER-to-Golgi lysosome enzyme trafficking. Shown is a comparison between WT conditions and deficiency of CLN6.

Inefficient organelle exit and depletion of cargo proteins upon deficiency of components of their receptor systems are commonly observed in the mammalian cell. For example, ERGIC-53 promotes ER export of immunoglobulin IgM and coagulation factors V/VII in complex with ERp44 and MCFD2, respectively (40, 41). In cells lacking functional ERGIC-53, its cargo proteins reach their destination at a slower rate and in reduced amounts (42, 43). Similarly, exit of LyzC from the Golgi is delayed upon deletion of the LyzC sorting protein, Cab45 (39). Additional examples from recent studies include a variety of receptor systems and cargo proteins—Vti1a/b for Golgi export of Neuropeptide-Y (44), SURF4 for the secretion of PCSK9 (45), and p24 for ER export of GPI-anchored proteins (46). Bulk flow or lower-affinity binding to other cargo receptors can explain slower cargo transport (47).

Recent studies focused on activation of TFEB, a master regulator of lysosomal biogenesis and function (48, 49), indicate that lysosomal enhancement may counteract disease progression in animal models of lysosomal storage disorders, including a model of Batten disease (50, 51). Thus, the finding that defects in the EGRESS complex result in reduced amounts of enzymes at the lysosome identifies CLN6 and CLN8 diseases as candidate conditions for testing lysosomal enhancement therapy (52). Interestingly, among the enzymes that are depleted upon CLN6 or CLN8 deficiency are TPP1 and CTSD, which are the defective proteins in two other NCL subtypes—CLN2 and CLN10, respectively (53–55). Both TPP1 and CTSD are involved in the degradation of subunit c of mitochondrial ATP synthase (SCMAS) (56–58), and studies that have characterized the composition of the storage material in various NCL subtypes have determined that SCMAS accumulates not only in CLN2 and CLN10 but also in CLN6 and CLN8 diseases (59–61). Our results thus establish a framework to interpret the accumulation of SCMAS upon CLN6 and CLN8 deficiency as being caused by the depletion of the SCMAS degrading proteins, TPP1 and CTSD, at the lysosome. These observations also suggest therapeutic opportunities based on the possibility to provide the depleted enzymes exogenously via enzyme replacement therapy, an option that is being tested for several lysosomal enzymes including CLN2 but that is not available for transmembrane proteins such as CLN6 and CLN8 (62–65).

In summary, our findings address the molecular mechanism underlying Batten disease caused by loss of functional CLN6 and identify the EGRESS complex as the mediator of the recruitment of lysosomal enzymes in the ER for Golgi transfer. These findings uncover a previously unappreciated complexity of the early steps of lysosomal enzyme trafficking and shed light on the molecular pathogenesis of CLN6 disease, which can henceforth be interpreted as a disorder of impaired lysosomal biogenesis.

## METHODS

### Animal husbandry

*Cln6^nclf^* mice on C57/BL6J background (*Cln6^−/−^* mice) were a gift from Dr. Susan Cotman (Harvard Medical School) and are available from the Jackson Laboratory (stock no: 003605). The *Cln8^mnd^* mouse line (*Cln8^−/−^* line) was purchased from the Jackson Laboratory (stock no: 001612). *Cln6^−/−^* and *Cln8^−/−^* double KO mice were generated by crossing the single knock-out lines. Food and water were provided *ad libitum*. For subcellular fractionation and ERG experiments we used mice at 6 weeks of age. Only males were used for all analyses. Investigators were blinded to mouse genotype during data acquisition, and no randomization was necessary. No data were excluded from this study.

### Molecular biology

Supplemental Table 1 lists the antibodies used throughout the study. Supplemental Table 2 lists the oligos used for cloning, genome editing, and PRC. CLN6 and CLN8 were cloned into myc and YFP vectors (backbone: pcDNA3.1 from Invitrogen) by retrotranscription of RNAs from HeLa and HEK293-T cells using the QuantiTect Reverse Transcription kit (Qiagen) followed by PCR-mediated amplification and plasmid insertion with the in-Fusion cloning kit (Clontech). The CLN6ΔL2 construct was obtained by removing the codons for amino acids 155-222 using the Q5 Site-Directed Mutagenesis kit (New England Biolabs).

### Immunoblotting

Before harvesting, cells were rinsed in cold PBS and lysed using RIPA buffer (50 mM Tris HCl, pH 7.4, 1% NP40, 0.5% sodium deoxycholate, 0.1% SDS, 150 mM NaCl, 2 mM EDTA) or NP40 lysis buffer (50 mM Tris HCl, pH 7.5, 150 mM NaCl, 1% Np-40 (v/v), 10% glycerol) with protease (Roche) and phosphatase (Roche) inhibitors (1:100). Cells were left in lysis buffer on a nutator for 1 hour at 4°C. After 1 hour of lysis the solution was centrifuged at 13,000 rpm for 20 minutes and the supernatant was collected. Protein concentrations were measured with the bicinchoninic acid protein assay kit (Pierce, Rockford, IL), using bovine serum albumin as standard. Lysates were separated on a sodium dodecyl sulfate polyacrylamide gel electrophoresis (SDS-PAGE) and then transferred to polyvinyl difluoride (PVDF) membranes. Blots were incubated in blocking buffer (5% dried skimmed milk in Tris-buffered saline, pH 7.4, and 0.2% Tween 20, TBST) followed by overnight incubation with appropriate antibodies, which were diluted in blocking buffer. The following day, membranes were washed 3 times for 10 min each in TBST before incubation for 1 hour with secondary HRP antibodies diluted in blocking buffer. Detection was carried out with ECL Western blotting detection reagent (GE Healthcare). Images were detected with ImageQuant LAS 4000 (GE Healthcare) and quantified by Fiji analysis software.

### Generation of CLN6 knock-out and CLN8 knock-in cells

We used CRISPR–Cas9 genome editing to introduce a deletion in exon 2 of the *CLN6* gene in HEK-293T cells. In brief, we designed two complementary oligonucleotide couples (Supplemental Table 2) using the online CRISPR design tool (http://crispr.mit.edu) coding for a guide RNA upstream of a protospacer adjacent motif (PAM) site in exon 2 of *CLN6*. The two oligos were annealed and subsequently cloned into the pX458 plasmid (66), followed by Sanger sequencing of the insert to confirm the correct sequence. HEK-293T cells were then transfected with 2 µg plasmid and split into a 96-well plate by single-cell deposition. DNA was isolated from the expanded single colonies when confluent enough and used in PCRs using oligos to amplify exon 2 of *CLN6* with CloneAmp HiFi PCR Premix (Clontech) according to the manufacturer’s instructions; clones that were unable to produce an amplicon were subsequently Sanger sequenced, to confirm deletion of exon 2. We inserted a Myc tag (5’-GAACAAAAACTCATCTCAGAAGAGGATCTG-3’) just before the stop codon of endogenous *CLN8* in HEK-293T cells by combining CRISPR–Cas9 with single-stranded oligodeoxynucleotides (ssODNs) as described (67). We selected recombinant clones by single colony expansion followed by gDNA extraction and PCR using oligos to amplify exon 3 of *CLN8*. Sanger sequencing confirmed the correct insertion of the Myc tag in a clone that was subsequently used for the described experiments.

### Quantitative real-time PCR

Total RNA was extracted from control and *Cln6^nclf^* mouse embryonic fibroblasts, and from control and *CLN6^−/−^* HEK-293T cells using the RNEasy kit (Qiagen) according to the manufacturer’s instructions. RNA was quantified using the Nano-Drop 8000 (Thermo Fischer) followed by cDNA synthesis using QuantiTect Reverse Transcription kit (Qiagen). The primers for PCR with reverse transcription reactions are listed in Supplemental Table 2. Quantitative real-time PCR was performed by using iQ SYBR Green Supermix on the CFX96 Touch Real-Time Detection System (Bio-Rad Laboratories) with the following conditions: 95°C, 5 min; (95°C, 10 s; 60°C, 10 s; 72°C, 15 s) x 40. Analyses were conducted using CFX manager software (Bio-Rad) and the threshold cycle (CT) was extracted from the PCR amplification plot. Relative gene expression was determined using methods described previously, normalizing to *GAPDH* (for human genes) and *cyclophilin* (for mouse genes). The change in messenger RNA level of the genes was expressed as fold change.

### RUSH Cargo Sorting Assay

WT and *CLN6*^−/−^ cells were cultured in a 96-well plate pre-coated for 2 hr with Poly-D-Lysine to aid in cell adherence. Cells were transfected for 24 hr using plasmids expressing test (CTSD, PPT1, GALNS) and control (LyzC) proteins fused with eGFP and a streptavidin binding protein (SBP) (pIRESneo3-LyzC-SBP-eGFP, pIRESneo3-SS-SBP-eGFP-CTSD, pIRESneo3-SS-SBP-eGFP-PPT1, and pIRESneo3-SS-SBP-eGFP-GALNS). Cells were incubated with 40 µM d-Biotin (SUPELCO) in DMEM for 10 min or without d-Biotin as a control to confirm the retention of the reporter. Cells were then washed once in PBS and fixed in 4% PFA in PBS for 15 min and used in confocal microscopy.

### Confocal microscopy analysis

Cells were grown on glass coverslips in 24-well plates overnight prior to treatment. Cells were transfected using Polyplus reagent, incubated for 6 hrs, and left to incubate for additional 36 hrs after changing the culture media. At the end of the treatment, cells were rinsed with PBS and fixed with 4% paraformaldehyde (PFA) in PBS at room temperature (RT) for 15 min. Cells were then rinsed three times with PBS and blocked with blocking reagent (0.1% saponin, 10% goat serum in PBS) for 60 min at RT. After blocking, cells were washed twice with PBS, followed by incubation with appropriate primary antibody in blocking reagent overnight at 4 °C. Cells were then washed three times with PBS and later incubated with appropriate Alexa-Fluor conjugated secondary antibodies against primary host animal for 1 h at RT in the dark. Cells were then washed five times with PBS and coverslips were then mounted with VECTASHIELD mounting media containing DAPI (Vector Laboratories). Images were acquired with Zeiss 710 and Zeiss 880 confocal microscopes (Carl Zeiss, Inc., Oberkochen, Germany). For the RUSH analysis of protein subcellular trafficking, imaging was performed on a GE Healthcare DVLive epifluorescence image restoration microscope using an Olympus PlanApo 60x/1.42 NA objective and a 1.9k x 1.9x pco.EDGE sCMOS_5.5 camera with a 1900×1900 FOV. The filter sets used were: DAPI, FITC, TRITC, and CY5. For co-localization analysis, quantification was done with organellar markers, pictures were analyzed using Coloc2 and plot profile plugin using ROI manager in Fiji software.

### Immunoprecipitation assay

HEK293-T cells after 24 or 48 hrs of transfection were scraped off from Petri dishes in cold PBS. Cells were centrifuged at 2000 rpm for 5 min and the pellet was re-suspended in NP40 lysis buffer (50 mM Tris HCl, pH 7.5, 150 mM NaCl, 1% Np-40 (v/v), 10% glycerol) with protease (Roche) and phosphatase (Roche) inhibitors (1:100). Cells were lysed for 30 min at 4°C on a nutator. Upon lysis, equal amounts of protein lysates (500 µg) were incubated with the indicated primary antibodies for 3 hrs at 4°C. After antibody incubation, samples were further incubated for 2 hrs in 40 µl pre-cleared protein-G agarose beads (Roche). Beads with immunocomplexes were centrifuged at 3,000 × g and washed four times in lysis buffer with intermittent incubations, during each wash, for 6 min on the nutatotor at 4°C. After the fourth wash, beads were resuspended in Laemmli SDS sample buffer with β-Mercaptoethanol heated at 95°C for 10 min and then analyzed by immunoblotting. For experiments involving CLN8, samples were incubated at 37°C to avoid formation of CLN8 dimers and smears observed at higher temperatures which are hard to resolve on a SDS gel.

### Flow cytometry analysis

HeLa cells were transfected with 200 ng YFP1- and 200 ng YFP2-tagged constructs, in combination with 200 ng of Ruby plasmid that was used as a reference for transfection efficiency. Cells were collected 48 hrs after transfection in PBS with 10% FBS. Fluorescence of 10,000 cells per sample was determined by flow cytometry using the BD LSRFortessa™ Cell Analyzer (BD Biosciences) with the HTS auto-sampler device. Only cells positive for YFP and Ruby were considered for the subsequent analysis of reconstituted YFP signal quantification.

### Subcellular fractionation

Three mouse livers per sample were pooled and centrifuged in a discontinuous Nycodenz (Progen Biotechnik) density gradient as previously described (68), with modifications. Briefly, we homogenized tissues in an assay buffer (0.25 M sucrose, pH 7.2) and centrifuged at 4,800 × g for 5 min, and then at 17,000 × g for 10 min. The sediment of the second centrifugation was washed at 17,000 × g for 10 min. Consequently it was resuspended 1:1 vol/vol in 84.5% Nycodenz, and placed on the bottom of an Ultraclear (Beckman) tube. Above this, a discontinuous gradient of Nycodenz was constructed using the following percentages from bottom to top: 32.8%, 26.3%, and 19.8% Nycodenz. Samples were then centrifuged for 1 h in an SW 40 Ti rotor (Beckman) at 141,000 × g. Lysosome-enriched fractions were collected from the 26.3/19.8 interface and diluted in 5-10 volumes of assay buffer. Finally, they were centrifuged at 37,000 × g for 15 min. Pellets were then resuspended in 500 µl of assay buffer.

### Enzyme assays

Enzyme activity assays were conducted by using fluorophore analogues of enzymes substrates, as previously described: TPP1 (69); GAA (70); GBA (71); GLB1 (72); NAGLU (73).

### Electroretinography (ERG)

Mice were dark-adapted overnight and then anesthetized by a single intraperitoneal injection of 22 mg/kg ketamine, 4.4 mg/kg xylazine and 0.37 mg/kg acepromazine. Pupils were dilated with a drop of tropicamide (1.0%) and phenylephrine (2.5%) and then corneas were anesthetized with a drop of proparacaine (1.0%). After 1 min, excess fluid was removed and a drop of hypromellose (2%) was placed on each cornea to keep it moistened and provide a good contact between the cornea and the ERG electrode (N1530NNC, LKC Technologies). All tests were performed under a dim red light and a feedback controlled heating pad was used to keep treated mice at a constant body temperature of 37°C. ERG recordings were performed using UTAS Visual Diagnostic System and EMWIN software (LKC Technologies, Gaithersburg, MD, USA). Scotopic ERGs were performed using six flash intensities (−34, −24, −14, −4, 0 and 10dB). Photopic ERGs were subsequently recorded. Mice were light-adapted to a 30 cd*s/m^2^ white background for 2 min after scotopic ERG recordings, and then photopic ERGs were recorded with flash intensities of 0, 10, and 25 dB. ERG data was plotted using GraphPad Prism5 software (GraphPad Software, La Jolla, CA, USA). All mice were analyzed at four weeks of age.

### CBM assay

HeLa cells were plated on coverslips, 24 hours before the experiment in 24-well plates in DMEM at 37°C in the presence of 5% CO_2_. Cells were transfected the next day with either myc-tagged CLN6 or myc-tagged CLN8. 24 hours after transfection the media was removed, cells were washed with PBS, and 1,3-Cyclohexanebis(methylamine) (CBM) was added to each well at a final concentration of 2 mM for a 90 min period were indicated.

### In vitro COPII vesicle budding reaction

Rat liver cytosol was prepared from fresh livers as previously described (31). HeLa cells were co-transfected with CLN6-myc and CLN8-myc, permeabilized as previously described (74), and used as donor membranes. Vesicle formation and purification was performed as described previously (74). In brief, donor membranes were incubated where indicated with rat liver cytosol (4 mg/ml), ATP regeneration system, 0.3 mM GTP, and Sar1A H79G (10 ng/µl) in a final volume of 100 µl. The reaction was carried out at 30°C for 1 h and terminated by incubation on ice for 5 min. Donor membranes were removed by centrifugation at 14,000 × g for 12 min at 4°C. The resulting supernatant was centrifuged at 115,000 × g for 25 min at 4°C to collect COPII vesicles. Vesicle pellets were then resuspended in 15 µl of buffer C (10 mM Tris-HCl, pH 7.6, 100 mM NaCl, 1% Triton X-100) and Laemmli sample buffer and heated at 55°C for 15 min. Vesicle samples and original donor membrane samples were resolved by SDS–PAGE, transferred onto polyvinylidene fluoride (PVDF) membranes, and subjected to immunoblotting to detect various ER-resident proteins and COPII cargo proteins.

### Sequence analysis

CLN6 sequence analysis was performed by retrieving CLN6 sequences from the NCBI database (https://www.ncbi.nlm.nih.gov/gene) or from the UCSC Genome Browser (http://genome.ucsc.edu) by BLAT searches using available CLN6 protein or DNA sequences. Local evolutionary rates of CLN6 protein sequence were calculated using the webtool Aminode (www.aminode.org) (37) by feeding the indicated sequences as inputs for the custom analysis. Downloaded data were graphically combined with a multiple protein alignment obtain with MULTALIN (http://multalin.toulouse.inra.fr/multalin) to generate the graph of the local evolutionary rates.

### Study approval

All animal studies were reviewed and approved of by the Institutional Animal Care and Use Committee of Baylor College of Medicine.

## AUTHOR CONTRIBUTIONS

M.S. conceived and supervised the study. L.B. and M.S. designed the experiments with the contribution of A.d.R., P.Z., A.E., R.P., J.S., D.R., J.C., R.N.S., S.Y.J, R.C. and R.W.S. L.B. performed most cell biology, molecular biology and mouse analyses. A.d.R., R.P. and J.S. performed molecular analyses. L.B. and P.Z. performed in vitro budding assay under the supervision of R.W.S. A.E., D.R. and L.B. performed ERG analysis under the supervision of R.C. L.B. and M.S. wrote the manuscript with input from all authors.

## ACKNOWLEDGMENTS

We thank H. Bellen, H. Zoghbi, M. Wang and T. Eissa for helpful discussion and K. Venkatachalam and H. Jafar-Nejad for critical reading of the manuscript. We thank Dr. Susan Cotman for the generous gift of mice from her *Cln6^nclf^* colony. We thank J. Von Blume for providing the pIRESneo3-LyzC-SBP-eGFP and pIRESneo3-SS-SBP-eGFP-CTSD plasmids. This work was supported by NIH grants NS079618 and GM127492 (to M.S.) and grants from the Gwenyth Gray Foundation, Beyond Batten Disease Foundation, and NCL-Stiftung (to M.S). This project was supported in part by IDDRC grant number 1U54 HD083092 from the Eunice Kennedy Shriver National Institute of Child Health and Human Development (Core: Integrated Microscopy). Imaging for this project was supported by the Integrated Microscopy Core at Baylor College of Medicine with funding from NIH (DK56338, and CA125123), CPRIT (RP150578, RP170719), the Dan L. Duncan Comprehensive Cancer Center, and the John S. Dunn Gulf Coast Consortium for Chemical Genomics.

## Notes

The authors have declared that no conflict of interest exists.

