## Supplemental Figures and Tables for "A CLN6-CLN8 complex recruits lysosomal enzymes at the ER for Golgi transfer"

### Supplemental Material:

**Supplemental Figure 1.** Depletion of lysosomal enzymes upon CLN6 deficiency.

**Supplemental Figure 2.** Interaction and localization assays for CLN6 and CLN8.

**Supplemental Figure 3.** Generation and validation of CLN6–/– cells.

**Supplemental Figure 4.** Evolutionarily constrained region analysis of CLN6.

**Supplemental Figure 5.** Generation and expression of the CLN6ΔL2 construct and tests for CLN6 dimerization.

**Supplemental Table 1.** List of antibodies.

**Supplemental Table 2.** Oligonucleotides used for cloning, gene/genome editing, and qRT-PC.

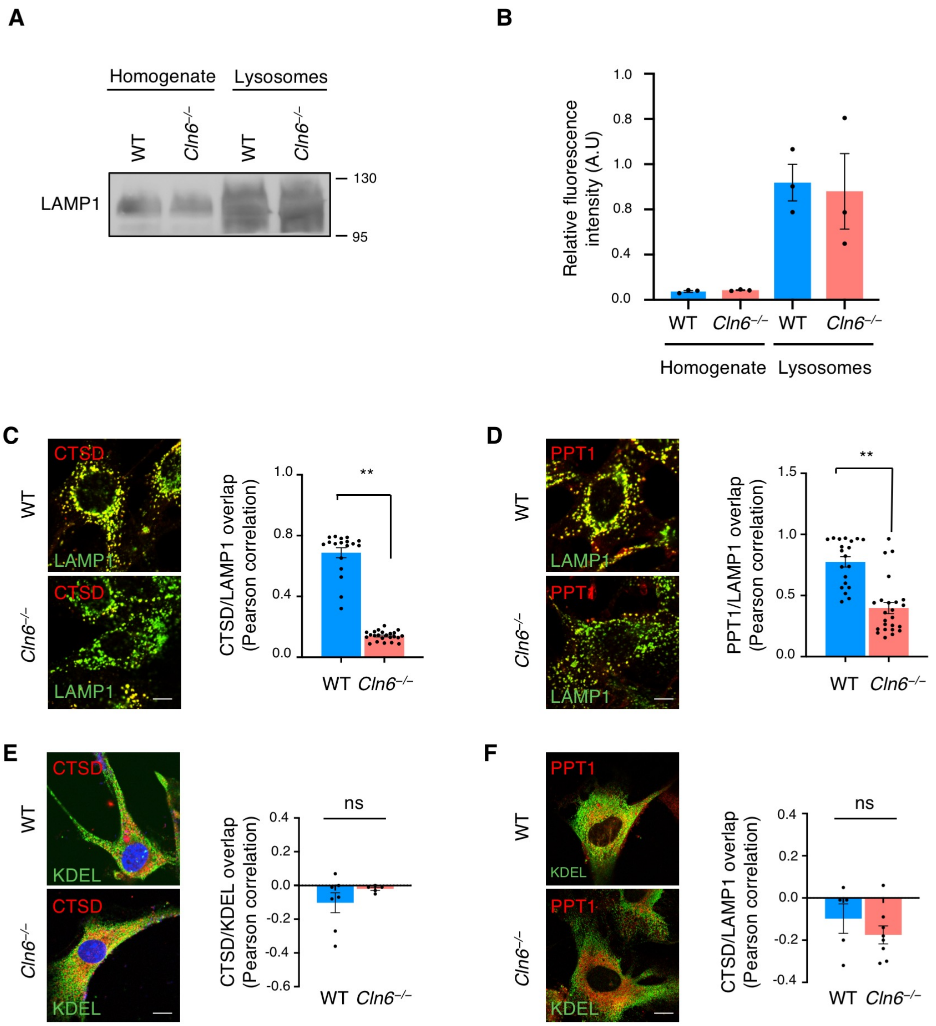


**Supplemental Figure 1. Depletion of lysosomal enzymes upon CLN6 deficiency. (A)** LAMP1 immunoblot showing lysosomal enrichment in collected fractions from WT and *Cln6*$^{-/-}$ samples. **(B)** HEX enzyme activity assay showing significant enrichment in the collected fractions from WT and *Cln6*$^{-/-}$ mice. (**C-F**) Confocal microscopy analysis of WT and *Cln6*$^{-/-}$ MEFs showing decreased overlap of the signals of endogenous CTSD and PPT1 with the lysosomal marker LAMP1 in *Cln6*$^{-/-}$ cells (**C** and **D**) and no changes in the overlap of CTSD and PPT1 with the ER marker KDEL (**E** and **F**). Data are means ± SEM ($n$ = 10, **P < 0.01, ns, not significant).

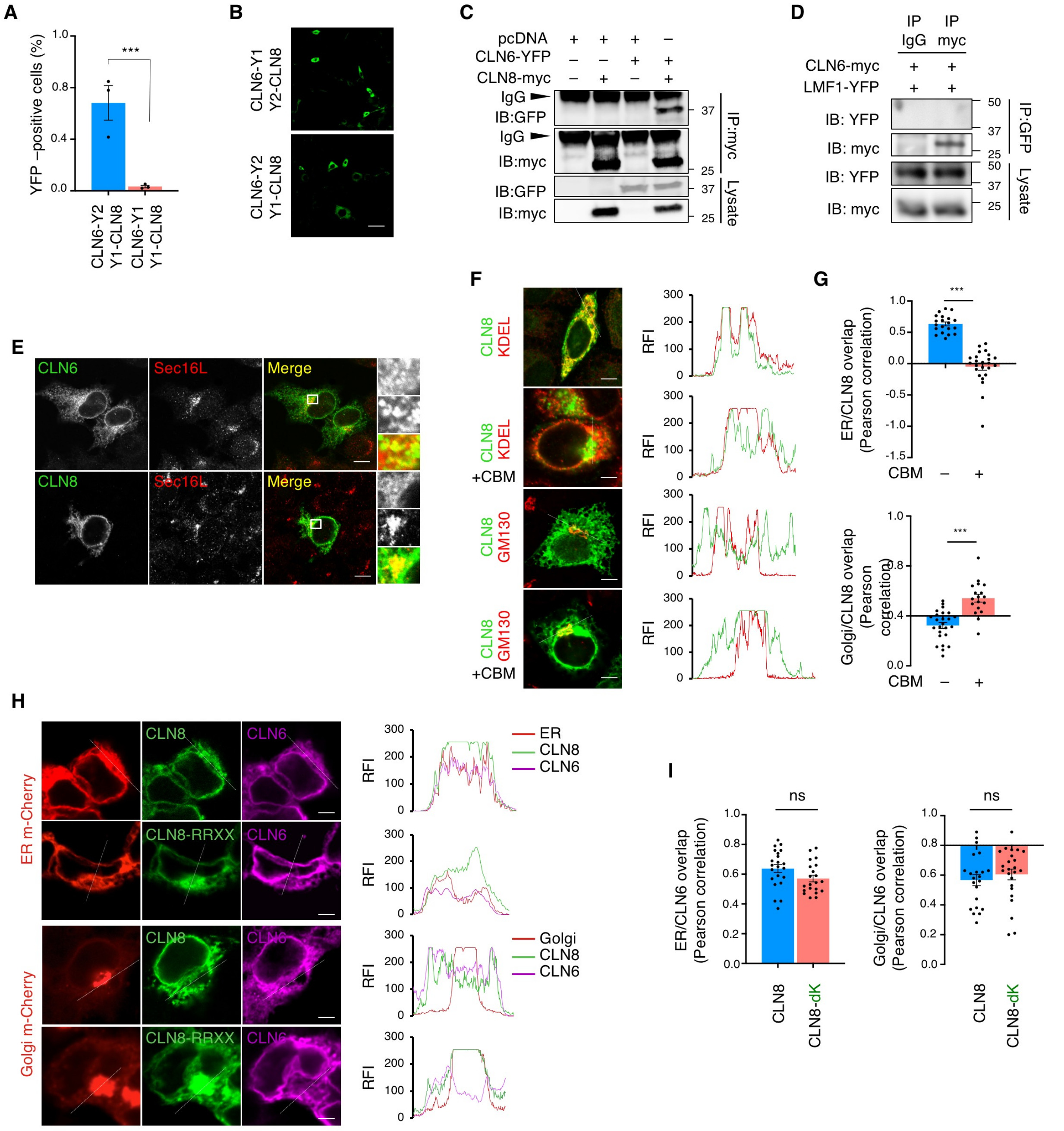

**Supplemental Figure 2. Interaction and localization assays for CLN6 and CLN8. (A)** BiFC assay of CLN6-Y2 with Y1-CLN8 in HeLa cells followed by flow cytometry. (**B**) BiFC assay of CLN6-Y2 with Y1-CLN8 and CLN6-Y1 with Y2-CLN8 in MEFs followed by confocal microscopy. Green signals represent the reconstituted YFP. Scale bar: 200 µm. (**C-D**) Co-IP analysis of CLN6 with CLN8 (**C**) and LMF1 (**D**). All proteins were tagged as indicated and transiently expressed. Co-IP assays were performed using the indicated antibodies and IgG as a negative control. Lysate represents 10% of the total cell extract used for IP. **(E)** Confocal microscopy showing that CLN6 and CLN8 colocalize with the ER exit site marker Sec16L (red). Scale bar: 20 µm. Inset magnifications (5x) are reported. (**F**) Confocal microscopy showing the subcellular localization of CLN8 upon treatment with CBM. Trace outline is used for line-scan analysis of Relative Fluorescence Intensity (RFI) of CLN8, GM130 (Golgi marker) and KDEL (ER marker) signals. Scale bar: 10 µm. (**F**) Pearson correlation analysis of the co-localization extent of full-length CLN8 and the ER marker KDEL or the Golgi marker GM130, with and without treatment with CBM. *n* > 10 images/experiment. **(H)** Confocal microscopy analysis showing ER localization of full-length CLN6 and Golgi localization of CLN8-RRXX (CLN8-dK) upon cotransfection of the two constructs in *CLN6*$^{-/-}$ cells. Trace outline is used for line-scan analysis of Relative Fluorescence Intensity of CLN8, CLN6, Golgi m-Cherry and ER m-Cherry signals. Scale bar: 10 µm. (**I**) Pearson correlation analysis of the co-localization extent of CLN6 with ER and Golgi markers when cotransfected with CLN8-dK in *CLN6*$^{-/-}$ cells. *n* > 10 images/experiment.

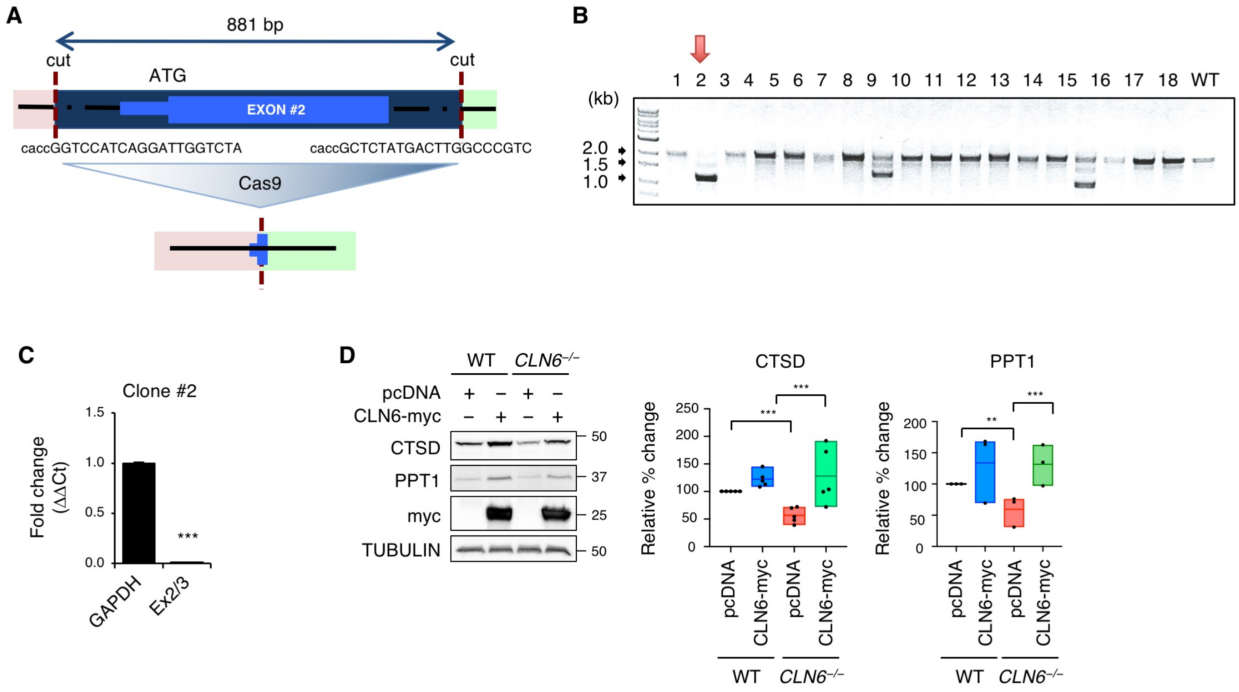


**Supplemental Figure 3. Generation and validation of *CLN6*$^{-/-}$ cells. (A)** Cartoon depicting the region from *CLN6* gene that was deleted to generate *CLN6*$^{-/-}$ cells by CRISPR/Cas9 genome editing. **(B)** PCR clone screening of individual HEK293-T cell lines to identify clones carrying deletion of *CLN6* exon 2 (CLN6$^{-/-}$) upon CRISPR/Cas9 genome editing. Expected WT amplification band: 2036 bp; expected *CLN6*$^{-/-}$ amplification band: 1155 bp. **(C)** qRT-PCR using RNA extracted from *CLN6*$^{-/-}$ clone #2 shows lack of expression of *CLN6* upon removal of exon 2 and flanking regions for a total of 881 bp. **(D)** Immunoblot analysis of WT and *CLN6*$^{-/-}$ cells showing decreased signal for endogenous CTSD and PPT1 in *CLN6*$^{-/-}$ cells, which was rescued by re-expression of myc-CLN6. Data are means ± SEM (*n* = 3, ***P < 0.001).

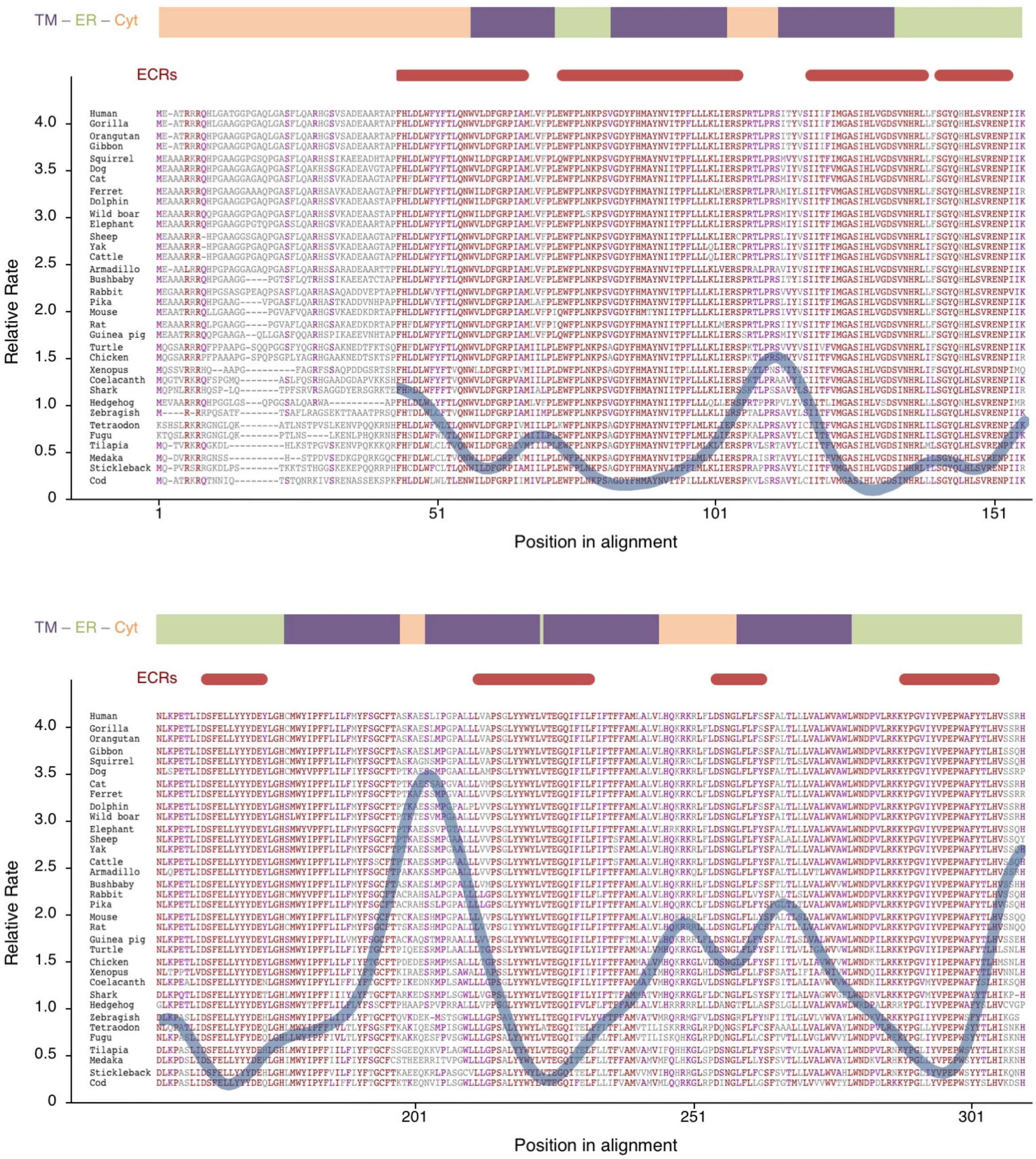


**Supplemental Figure 4. Evolutionarily constrained region analysis of CLN6.** Shown is a multi-alignment of CLN6 protein sequences along with a plot of local evolutionary rates. The red plot reports the average number of amino acid substitutions per site, taking into account the evolutionary relationships among proteins. Indicated are the protein transmembrane domains (TM, purple lines), cytosolic domains (Cyt, orange lines) and luminal domains (ER, green lines). Evolutionarily constrained regions (ECRs) are indicated with red lines.

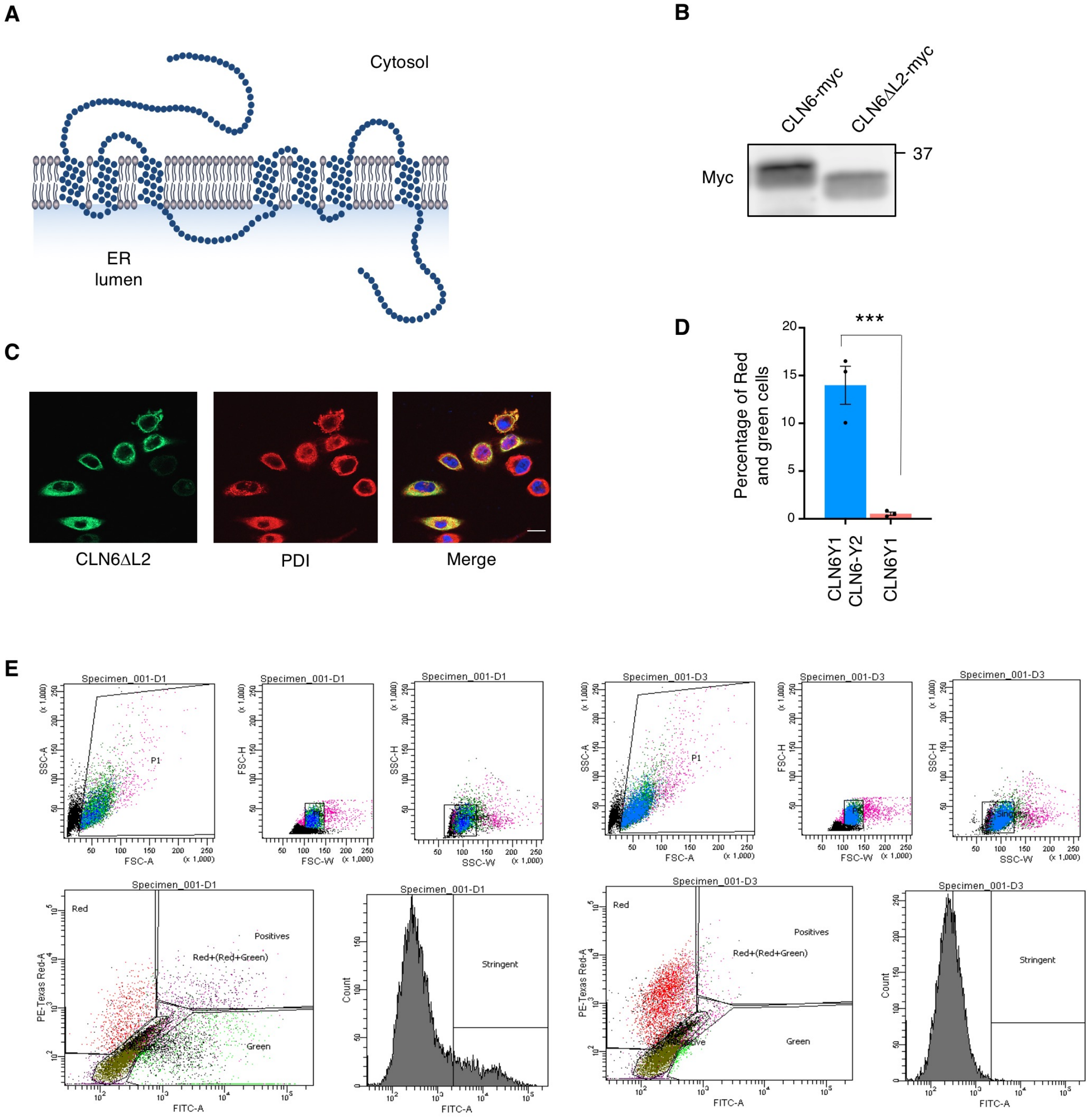


**Supplemental Figure 5. Generation and expression of the CLN6ΔL2 construct and tests for CLN6 dimerization. (A)** Cartoon structure of CLN6ΔL2. **(B)** Protein expression of full-length CLN6 and CLN6ΔL2, both tagged with myc. **(C)** Confocal microscopy analysis showing co-localization of CLN6ΔL2 (green) with the ER marker PDI (red). Scale bar: 20 μm. **(D)** Quantification of BiFC of CLN6-Y1 with CLN6-Y2 in HEK-293T cells by flow cytometry. CLN6-Y1 is used as a negative control. **(E)** Representative gating for BiFC-flow cytometry analysis. Cells were co-transfected with split-YFP vectors and a Ruby plasmid (red fluorescence) that was used to normalize BiFC readings for transfection efficiency. CLN6 proteins form homodimers and we used this property to set up the gates for BiFC-flow cytometry assay. Shown are examples of histograms of BiFC data for CLN6-Y1/CLN6-Y2 *(left)* and CLN6-Y1 (control, *right*).

**Supplemental Table 1. List of antibodies.**

| Antibodies | Supplier | Catalog # | Host | Working dilution |
|---|---|---|---|---|
| | | *Primary antibodies* | | |
| anti-C-myc | Sigma-Aldrich | C3956-2MG | Rabbit | 1:1000 |
| anti-CTSB (CA10) | Millipore | IM27L-100UG | Mouse | 1:1000 |
| anti-CTSD (R-20) | Santacruz | sc-6487 | goat | 1:1000 |
| anti-CTSD (CTD-19) | Abcam | ab6313 | Mouse | 1:1000 |
| anti-GALNS | Abcam | ab187516 | goat | 1:1000 |
| anti-GAPDH (6C5) | Santacruz | sc-32233 | Rabbit | 1:1000 |
| anti-GFP | Abcam | ab13970 | Chicken | 1:1000 |
| anti-GFP | Cell Signaling | 2956S | Rabbit | 1:1000 |
| anti-GM130 | Abcam | ab526449 | Rabbit | 1:200 |
| anti-KDEL (10C3) | Abcam | ab12223 | Mouse | 1:200 |
| anti-LAMP1 (1D4B) | Santacruz | sc-19992 | Rat | 1:250 |
| anti-LAMP2 (H4B4) | Santacruz | sc-18822 | Mouse | 1:1000 |
| Anti-PPT1 | Santacruz | sc-130726 | Rabbit | 1:500 |
| Anti-PPT1 | Sigma | HPA021546-100UL | Rabbit | 1:1000 |
| Anti-TPP1 | Santacruz | sc-393961 | Mouse | 1:1000 |
| Anti-TPP1 (H-300) | Santacruz | sc-66836 | Rabbit | 1:500 |
| Anti-NAGLU | Abcam | ab72178 | Rabbit | 1:500 |
| | | *Secondary antibodies* | | |
| Alexa Fluor® 488 Anti-Rat IgG | Invitrogen | A-11006 | Goat | 1:1000 |
| Alexa Fluor® 488 Anti-Chicken IgG | Invitrogen | A-11039 | Goat | 1:1000 |
| Alexa Fluor® 488 Anti-Mouse IgG | Invitrogen | A-21200 | Chicken | 1:1000 |
| Alexa Fluor® 594 Anti-Mouse IgG | Invitrogen | A-21201 | Chicken | 1:1000 |
| Alexa Fluor® 594 Anti-Rabbit IgG | Invitrogen | A-21202 | Donkey | 1:1000 |
| Alexa Fluor® 633 Anti-Mouse IgG | Invitrogen | A-21052 | Goat | 1:1000 |
| Alexa Fluor® 633 Anti-Goat IgG | Invitrogen | A-21082 | Donkey | 1:1000 |
| ECL anti-rabbit IgG-HRP | GE Healthcare | NA9340-1ML | Donkey | 1:5000 |
| ECL anti-chicken IgY-HRP | Santacruz | sc-2428 | Goat | 1:5000 |
| ECL anti-goat IgG-HRP | Santacruz | sc-2020 | Donkey | 1:5000 |
| ECL anti-rat IgG-HRP | GE Healthcare | NA9350-1ML | Goat | 1:5000 |
| ECL anti-mouse IgG-HRP | GE Healthcare | NA9310-1ML | Sheep | 1:5000 |

**Supplemental Table 2. Oligonucleotides used for cloning, gene/genome editing, and qRT-PC.**

| Name | Forward Primer | Reverse Primer |
|---|---|---|
| | *Cloning and mutagenesis* | |
| pIRESneo3-SS-SBP-eGFP-PPT1 | agctgtacaagggccgggacccgccggcgccgc | tgatcagttatctagttatccaaggaatggtatgatgtgg |
| pIRESneo3-SS-SBP-eGFP-GALNS | agctgtacaagggccgggccccgcagccccccaacat | tgatcagttatctagttagtgggaccagaggcacttctt |
| CLN6 infusion short | cagtgtggtggaattcatggaggcgacgcggagg | ccaccgccaccatcgatgtgccgactgctgacgtgaagg |
| CLN8ΔL | ctggtcagcagcctgtatc | agcttggagattgaccaaag |
| CLN8dK | ggccctctagactcgagctatggccttcgtcgacgcagcagctgcccgttgccttc | gaaggcaacgggcagctgctgcgtcgacgaaggccatagctcgagtctagagggcc |
| | *Genome editing* | |
| CRISPR_CLN6_1 | caccggtccatcaggattggtcta | caccgctctatgacttggcccgtc |
| CRISPR_CLN6_2 | aaactagaccaatcctgatggacc | aaacgacgggccaagtcatagagc |
| CRISPR_CLN8_B | caccgtgtagggcccggcccgtgt | aaacacacgggccgggccctacac |
| CRISPR_CLN8_D | caccgcgccctttacctgcgtttcc | aaacggaaacgcaggtaaagggcgc |
| DNA_check_CLN6 | ttgggaaaagcttcatcagg | ggtgctccacttcctttcct |
| qRT_CRISPR CLN6 | aggccaggcatggctct | ggaataccagcatggcaatg |
| CLN8_entryclone | ttgctcgagggaattcatgaatcctgcgagcgatggg | cgcaccgggcgaattccttacatattcagatcctcttctga |
| CLN8_KI_check | agcataggaccgtgtgttcc | aaaggtagccaggtgtggtg |
| | *RT-qPCR* | |
| *Cyclophilin* | ggcaaatgctggaccaaacacaa | gtaaaatgcccgcaagtcaaaag |
| *Ctsb* | ttgcgttcggtgaggacatag | aaatgcccaacaagagccg |
| *Ctsd* | cgtcctttgacatccactacgg | tggaaccgatacagtgtcctgg |
| *Gaa* | ctcctacccaggtcc | atggccaggctcttgttgtcag |
| *Glb1* | aaatggctggcagtccttctg | acctgcacggttatgatcggt |
| *Tpp1* | cccctcatgtggattttgtgg | tggttctggacgttgtcttgg |
